# Exploring the neuroimmune cellular landscape in the skin of subjects with fibromyalgia

**DOI:** 10.1101/2025.11.05.686713

**Authors:** Carlos E. Morado-Urbina, Kirill Agashkov, Matthew Hunt, Macarena Tejos-Bravo, Karolina af Ekenstam, Silvia Fanton, Sigita Venckute-Larsson, Katalin Sandor, Emilie Linderoth, Alexandra Kuliszkiewicz, Theodor Arlestig, Monika Löfgren, Eva Kosek, Camilla I. Svensson

## Abstract

Fibromyalgia (FM) is a chronic disorder involving widespread pain, fatigue, and cognitive impairment. Although its pathogenesis remains uncertain, FM patients exhibit hypersensitivity, reduced intraepidermal nerve fiber density (IENFD), and more recently shown, skin immune dysregulation, possibly manifesting in cutaneous alterations in various cell populations. To characterize the cutaneous cellular landscape in FM, we assessed the sensory profile and skin cell populations across different layers within the same cohort. FM patients, compared to healthy controls (HC), showed lower pain thresholds across multiple body areas, with no differences in thermal detection. While our findings confirm previous reports for reduced PGP9.5+ IENF and increased density in mast cells in FM, we also identified novel changes, particularly in the dermis. We observed elongated thinly myelinated NF200+ fibers and reduced density in non-nerve-associated S100B+ Schwann cells in FM compared to HC. Notably, dermal CD68+ and CD163+ populations were significantly reduced in FM, accompanied by morphological changes. The CD163+ population correlated negatively and significantly against IENFD. These findings suggest that, beyond intraepidermal nerve loss, FM involves broader neuroimmune alterations in the skin, particularly within the dermis, offering new insights into its pathophysiology and establishing a foundation for future studies exploring the functional implications of these changes.

**SUMMARY:** Among the alterations observed in patients with fibromyalgia (FM) are changes in skin innervation and differences in the density of certain immune cell populations. To better characterize FM from an integral perspective, we examined both the sensory profile and the cutaneous cellular landscape of FM patients in comparison with healthy controls (HC). FM patients exhibited lower pain thresholds across multiple body areas, with no differences in thermal detection. Consistent with previous findings, FM skin biopsies revealed reduced intraepidermal nerve fiber density and increased mast cell numbers. Beyond these known alterations, our study identified novel dermal changes, including elongated thinly myelinated NF200+ fibers, reduced density of non–nerve-associated S100B+ Schwann cells, and decreased CD68+ and CD163+ macrophage populations exhibiting morphological alterations. Notably, CD163+ cell density correlated negatively with intraepidermal nerve fiber density, highlighting potential neuroimmune mechanisms underlying the pathophysiology of FM.

## INTRODUCTION

Fibromyalgia (FM) is a multifaceted chronic pain disorder that affects a significant portion of the global population, with a mean estimated global prevalence of 2.7% (1). This condition is characterized by a complex array of symptoms, including widespread pain, heightened sensitivity to mechanical and cold stimuli, physical and mental fatigue, sleep disturbances, and cognitive impairments (2–4). Despite its prevalence and substantial impact on the quality of life, the pathogenesis of FM remains enigmatic, and its diagnosis and management are challenging. Although FM pain is severe and widespread, it is not associated with identifiable injuries in peripheral tissues (5). Traditionally, FM has been attributed to persistent alterations in nociceptive processing. While substantial evidence supports CNS changes in FM (5–7), emerging research increasingly highlights also abnormalities in peripheral sensory neurons that innervate the skin, particularly within the epidermis. These abnormalities include spontaneous activity, heightened excitability, and decreased intraepidermal nerve fiber density (IENFD) (8–12).

Interestingly, individuals with FM not only experience altered cutaneous sensitivity, but also frequently report other skin-related symptoms, such as chronic itch, hyperhidrosis, and burning sensations (13–15). Moreover, a higher prevalence of FM has been reported in patients with skin disorders and vice versa (15). For instance, patients with chronic urticaria (CU) exhibit a significantly higher incidence of FM compared to controls (26% vs. 8%) (15, 16). Similarly, an increased prevalence of FM has been documented in patients with acne vulgaris (22% vs. 5%) and vitiligo (34% vs. 11%) compared to healthy individuals (17, 18). Collectively, these findings underscore the intricate relationship between FM and skin manifestations, highlighting the importance of investigating the cutaneous tissues in the FM pathology.

It is well established that the activation of skin resident immune cells can sensitize and activate nociceptors through local release of proinflammatory and algogenic factors (19, 20). Communication also occurs between neurons and cells like keratinocytes in the epidermis and fibroblasts in the dermis. These cells can release various signaling molecules, including ATP, neurotrophic factors, axonal guidance and repellent molecules, which influence neuronal development, activity, and function (21). Schwann cells may be involved in orchestrating local immune responses and, like keratinocytes, can also modulate pain sensation (22–24). Additionally, changes in the density of Langerhans cells, one of the main resident immune cells in the epidermis, have recently been linked to the development of mechanical allodynia in models of painful diabetic neuropathy (25). Overall, these findings emphasize the critical role of neuroimmune interactions in the skin, where changes in sensory neurons and immune cells could both contribute to pathophysiological mechanisms in chronic pain conditions.

Several studies have identified cellular changes in the skin of FM patients. The most commonly reported alteration is a reduction in epidermal innervation (8–10, 26). In addition, an increased number of mast cells has been noted in the FM skin (9, 27, 28), leading to the hypothesis of a low-grade inflammatory environment in FM (29). Notably, the presence of NK-cells near dermal nerve fibers has also been observed in FM, a finding rarely seen in healthy skin (30). Furthermore, studies have reported reduced activity of mitochondrial respiratory enzymes and a decline in bioenergetic capacity in the skin of FM patients compared to healthy controls (31). These alterations can disrupt cellular processes, including inflammation and stress responses, potentially exacerbating skin-related symptoms and contributing to peripheral nociceptor sensitization and enhanced pain sensation.

While independent research groups have made valuable contributions by uncovering various skin alterations in FM, a comprehensive assessment of the sensory profile and the cutaneous cellular landscape, integrating both the epidermis and dermis, is essential for a more complete understanding of the condition. The aim of this study was to assess the presence of immune cells and nerve fibers in the skin of a cohort of FM patients who had undergone detailed sensory profiling.

## RESULTS

### Characterization of the study cohort

A total of 16 female subjects with FM (42 ± 11.7 years) and 16 healthy controls (HC, 54.8 ± 8.1 years) were included in the study. All FM subjects were screened by a specialist to confirm compliance with the ACR1990 and ACR2016 classification criteria (2, 3). Demographic and clinical data for study subjects are shown in Table 1. Age was lower in FM subjects compared with the control group (p = 0.0038), while BMI was higher in FM subjects (27.6 ± 2.9) than in HC (23.8 ± 2.7, p = 0.0004). As expected, FM subjects reported significantly elevated pain levels at the time of their visit compared to HC (62.9 ± 19.2 vs. 0.9 ± 1.8, p < 0.0001), as well as during the week prior to the visit (73.8 ± 11.8 versus 3.3 ± 2.2 in HC, p < 0.0001). The mean FIQ scores were higher in the FM group (70.7 ± 9.2) compared to the HC group (3 ± 2.3, p< 0.0001).

**Table 1.**
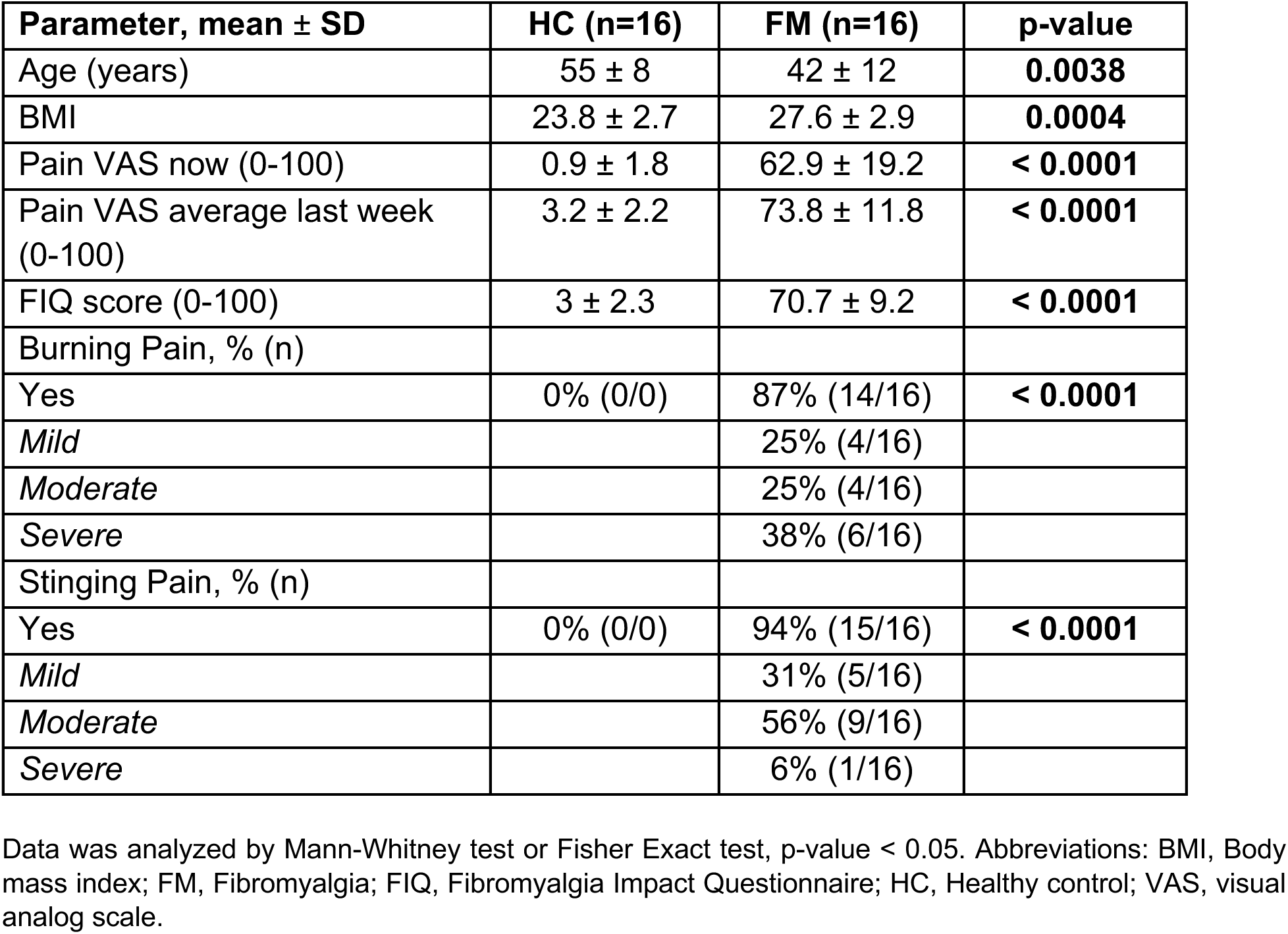
Demographic and clinical characteristics of study population.

| Parameter, mean ± SD | HC (n=16) | FM (n=16) | p-value |
| --- | --- | --- | --- |
| Age (years) | 55 ± 8 | 42 ± 12 | **0.0038** |
| BMI | 23.8 ± 2.7 | 27.6 ± 2.9 | **0.0004** |
| Pain VAS now (0-100) | 0.9 ± 1.8 | 62.9 ± 19.2 | **< 0.0001** |
| Pain VAS average last week (0-100) | 3.2 ± 2.2 | 73.8 ± 11.8 | **< 0.0001** |
| FIQ score (0-100) | 3 ± 2.3 | 70.7 ± 9.2 | **< 0.0001** |
| Burning Pain, % (n) |  |  |  |
| Yes | 0% (0/0) | 87% (14/16) | **< 0.0001** |
| *Mild* |  | 25% (4/16) |  |
| *Moderate* |  | 25% (4/16) |  |
| *Severe* |  | 38% (6/16) |  |
| Stinging Pain, % (n) |  |  |  |
| Yes | 0% (0/0) | 94% (15/16) | **< 0.0001** |
| *Mild* |  | 31% (5/16) |  |
| *Moderate* |  | 56% (9/16) |  |
| *Severe* |  | 6% (1/16) |  |

Data was analyzed by Mann-Whitney test or Fisher Exact test, p-value < 0.05. Abbreviations: BMI, Body mass index; FM, Fibromyalgia; FIQ, Fibromyalgia Impact Questionnaire; HC, Healthy control; VAS, visual analog scale.

### Widespread heightened thermal and mechanical pain sensitivity in FM with intact detection thresholds

Using the short form of the McGill pain questionnaire, we found that both burning and stinging pain were frequently reported by subjects with FM, affecting 88% (14/16) and 94% (15/16) of FM subjects, respectively. These pain features were not reported by any of the HC. As part of the sensory characterization, we used quantitative sensory testing (QST) in different body areas. We found similar detection thresholds for warmth and cold temperatures in FM and HC (Fig. 1). These results were observed for the warm detection threshold (WDT) and cold detection threshold (CDT), both in the arm (WDT: FM 34.6 ± 0.8 °C vs HC 35 ± 0.9 °C, p = 0.2; CDT: FM 30.8 ± 0.5 °C vs HC 31 ± 0.4 °C, p = 0.5, Fig. 1A, B) and the thigh (WDT: FM 34.8 ± 0.71 °C vs HC 35.1 ± 1.2 °C, p = 0.3; CDT: FM 30.3 ± 0.8 °C vs HC 30.1 ± 1.2 °C, p = 0.8, Fig. 1C, D).

**Figure 1.**
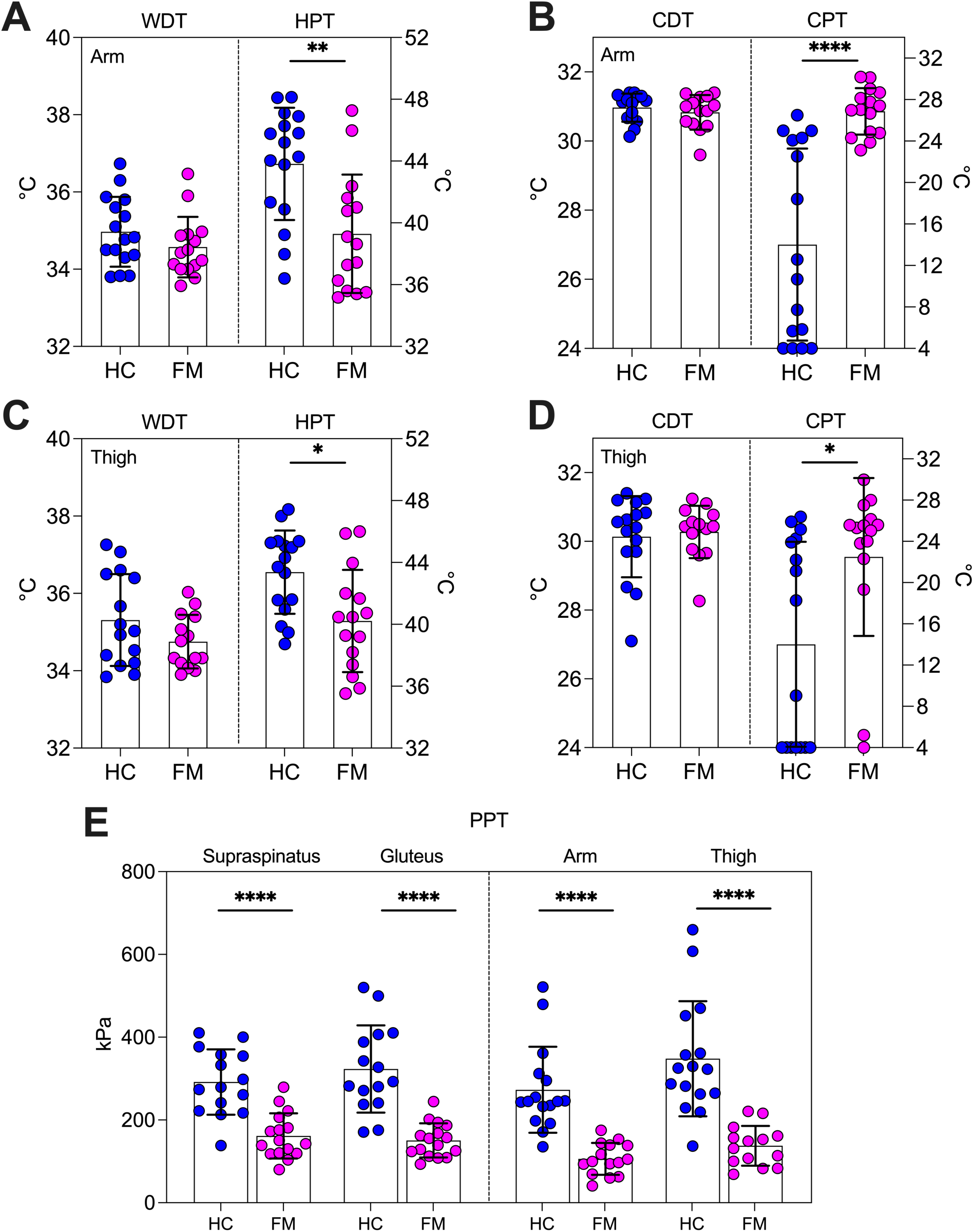
Sensory detection and pain thresholds across body sites in patients with FM. Thermal detection thresholds (left y-axis) and pain thresholds (right y-axis) were assessed in HC and patients with FM, both in the dorsal part of the arm (A, B) and the lateral side of the thigh (C, D). Patients with FM exhibited significantly increased pain sensitivity to both heat and cold, with no evidence of warm or cold hypoesthesia, as detection thresholds were similar to those of HC. Additionally, PPTs were measured bilaterally in the supraspinous and gluteus muscle as tender points according to ACR-1990 criteria (E, left panel). Additionally, PPTs in the arm and thigh were also measured (E, right panel), with FM patients showing significantly lower PPTs compared to HC at all sites. Data are expressed as mean ± SD and were analyzed using the Mann–Whitney test. *p < 0.05; **p< 0.01; ****p < 0.0001. HC N= 16, FM N=15-16. CDT, cold detection threshold; CPT, cold pain threshold; HPT, heat pain threshold; PPT, pressure pain threshold; WDT, warm detection threshold.

Compared to HC, heat and cold pain sensitivity was higher in FM (Fig. 1A-D). The significantly lower heat pain threshold (HPT) and cold pain thresholds (CPT) were observed in the arm (HPT: FM 39.3 ± 3.8 °C vs HC 43.8 ± 3.6 °C, p = 0.004; CPT: FM 26.9 ± 2.2 °C vs HC 14 ± 9.3 °C, p < 0.0001, Fig. 1A, B) and the thigh (HPT: FM 40.2 ± 3.3 °C vs HC 43.4 ± 2.7 °C, p = 0.015; CPT: FM 22.5 ± 7.7 °C vs HC 14 ± 9.9 °C, p = 0.007, Fig. 1C, D). A sensation of warmth on skin cooling after CPT testing (paradoxical heat sensation) was reported by 13% of FM patients (2/16) and 6% HC (1/16) (p = 1).

Additionally, FM patients showed a generalized increase in mechanical pain sensitivity reflected as lower pressure pain thresholds (PPTs) compared to HC (Fig. 1E). This mechanical hypersensitivity was found in tender points as the supraspinatus (FM 161.5 ± 54.5 kPa vs HC 291.7 ± 79 kPa, p < 0.0001), and the gluteus muscle (FM 150.7 ± 41.1 kPa vs HC 323.1 ± 105.3 kPa, p < 0.0001).

Furthermore, the arm (FM 106 ± 38.5 kPa vs HC 273 ± 104 kPa, p < 0.0001) and the thigh (FM 137.6 ± 48.1 kPa vs HC 347.9 kPa ± 138.9, p < 0.0001) also had decreased PPT in patients with FM.

### Patients with FM exhibit enhanced pain sensitivity during suprathreshold stimulation

To test whether sensitivity to suprathreshold pain was also different in FM, we determined the temperature corresponding to a pain rating of 4/10 (moderate-to-strong pain) or 7/10 (very strong pain), respectively, on a Borg10 scale (Fig. 2), and curves were plotted including the previously measured pain thresholds (HPT, CPT, PPT).

**Figure 2.**
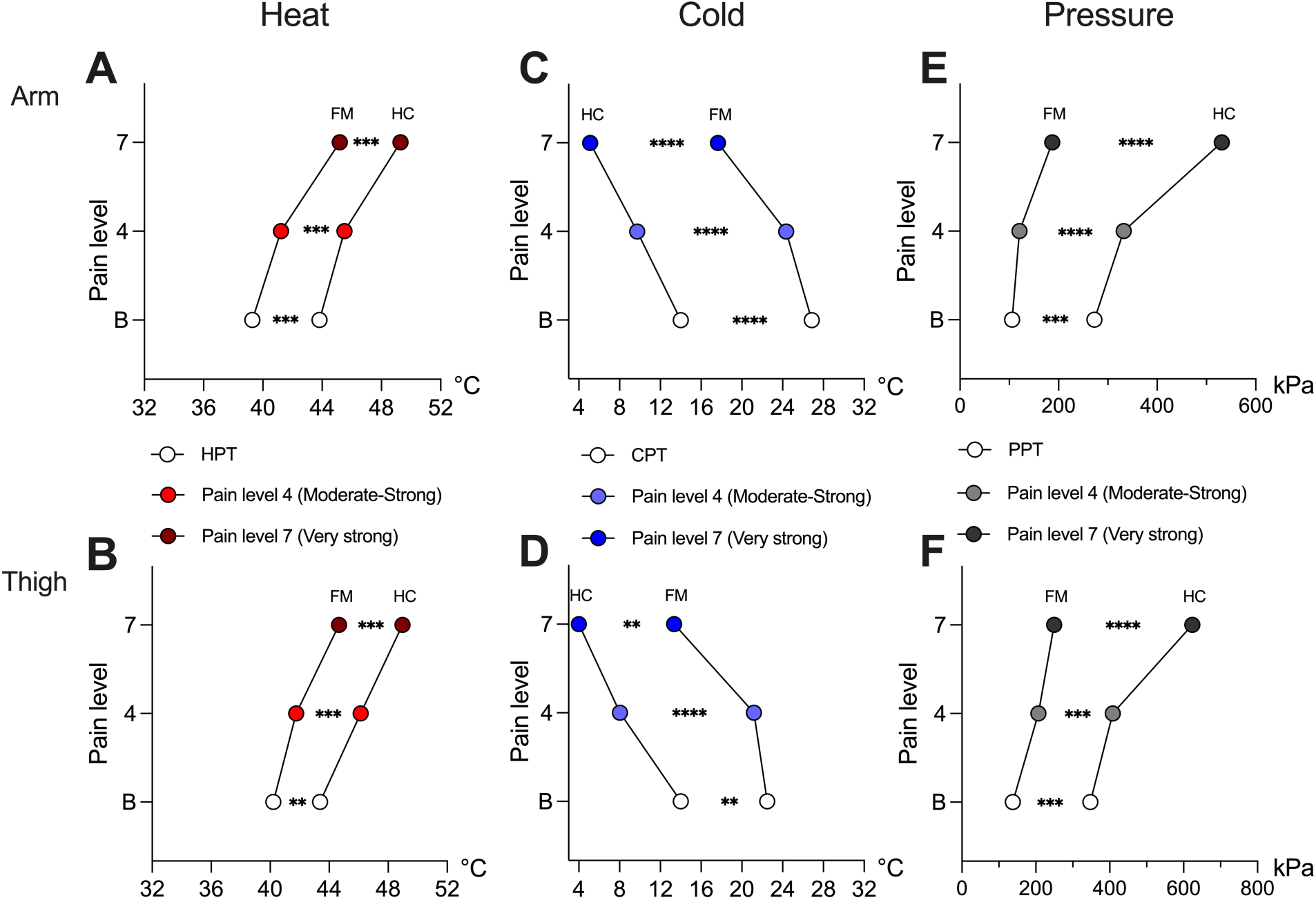
Suprathreshold pain sensitivity to temperature and pressure. Participants indicated when the applied temperature (°C) or pressure (kPa) reached a level of 4/10 (moderate-strong pain) or 7/10 (very strong pain) on a Borg scale. Stimuli were applied to the arm (A, C, E) and thigh (B, D, F) and data was integrated with pain thresholds (HPT, CPT or PPT). For heat, FM patients required significantly lower temperatures to reach 4/10 and 7/10 pain in both sites, with FM curves shifted to the left (A, B). For cold, FM patients reported a pain level of 4/10 and 7/10 at significantly warmer temperatures than HC in the arm (C) and thigh (D). The pressure required to elicit 4/10 and 7/10 levels of pain in both the arm and thigh (E, F) was significantly lower in FM compared to HC. Data are presented as mean ± SD. For the comparison of stimulation intensity for both pain levels across groups, a RM Two-way ANOVA test was used, **p< 0.01; ***p < 0.001; ****p < 0.000. HC N = 16, FM N = 15-16. CPT, cold pain threshold; HPT, heat pain threshold; PPT, pressure pain threshold.

Mean warm temperatures for pain ratings of 4/10 at the arm and thigh were significantly lower in FM (arm 41.2 ± 3.8°C; thigh 41.8 ± 3.9°C) compared to HC (arm 45.5 ± 2.6°C; thigh 46.1 ± 1.8°C, p < 0.001 for both areas). Similarly, temperatures to achieve a 7/10 level of pain were also lower in FM (arm 45.2 ± 2.9°C; thigh 44.7 ± 3.6°C) compared to HC (arm 49.3 ± 1°C; thigh 49 ± 1.1°C, p < 0.001 for both areas), both in the arm and the thigh (Fig. 2A, B), reflected on a clear shift to the left in FM temperatures curves.

Regarding cold temperatures, FM patients reported a pain intensity of 4/10 with temperatures in the range 27-17°, while HC reported pain with temperatures < 15°C, with significantly shifted temperatures curves. Beside changes in CPT, this was observed for stimulus corresponding to a 4/10 in both the arm (FM 24.3 ± 6.3°C vs HC 9.6 ± 8.6°C, p < 0.0001) and the thigh (FM 21.2 ± 9.1°C vs HC 7.8 ± 8.2°C, p < 0.0001) and replicated for an intensity of 7/10 in both areas (arm FM 17.7 ± 8.5°C vs HC 4.8 ± 2.7°C, p < 0.0001; thigh FM 13.3 ± 8.5°C vs HC 3.6 ± 0.5°C, p < 0.01) (Fig. 2C, D).

In the arm, 50% (8/16) of HC reached the lower limit of cold stimulation (4°C) during the first suprathreshold stimulation, and 75% (12/16) for the second stimulation. This was the case in 7% (1/15) and 13% (2/15) of FM patients for 4/10 and 7/10 level of pain, respectively. In the thigh, 73% (11/16) of HC reported a pain level of 4/10 at 4°C, while 100% (15/15) reported 7/10 of pain with 4°C. In FM patients, only 20% (3/15) reported 4/10 level of pain with 4°C, while 47% (7/15) reached 4°C for a 7/10 level of pain. This result reinforces a significantly heightened perceived intensity of cold in FM patients compared to HC.

Finally, the pressure at which FM patients reported 4/10 level of pain was significantly lower compared to HC, with FM curves shifted to the left for the arm and thigh. To achieve an intensity of 4/10, 120.5 ± 37.6 kPa were necessary in the arm and 207.1 ± 80.6 kPa in the thigh of FM subjects. For HC, significantly higher pressure was necessary in the arm (332.1 ± 94.4 kPa, p < 0.0001) and the thigh (408.5 ±138.2 kPa, p < 0.0001). In the case of a pain level of 7/10, FM subjects reported this intensity with 187 ± 75.6 kPa in the arm and 249.9 ± 104.9 in the thigh, while significantly more pressure was necessary in HC, with 531.3 ± 227.2 kPa in the arm and 623.8 ± 257.9 in the thigh (p < 0.0001 for both areas) (Fig. 2E, F), reinforcing the presence of profound mechanical hyperalgesia in FM.

### FM skin shows distinct differences in epidermal and dermal innervation compared to HC

In this well-characterized cohort of FM patients exhibiting the classic hyperalgesic phenotype, we quantified the density of various skin-cell populations. First, to evaluate innervation of epidermis and dermis, we employed two commonly used general neuronal markers: Protein gene product 9.5 (PGP9.5), a ubiquitin carboxyl-terminal hydrolase enzyme that is present in neurons and neuroendocrine cells, and tubulin beta-3 chain (TUBB3), which is expressed in microtubules within axons. The number of PGP9.5^+^ fibers crossing the dermal-epidermal border was used to quantify the IENFD (Fig. 3A). The results showed a significant reduction in the IENFD in FM skin (mean 6.8 ± SD 2.9 fibers/mm) compared to HC (10.1 ± 5.3 fibers/mm, p = 0.04) (Fig. 3B). The number of TUBB3^+^ crossings into the epidermis was also quantified (Fig. 3C); however, no significant differences were observed between the groups (Fig. 3D).

**Figure 3.**
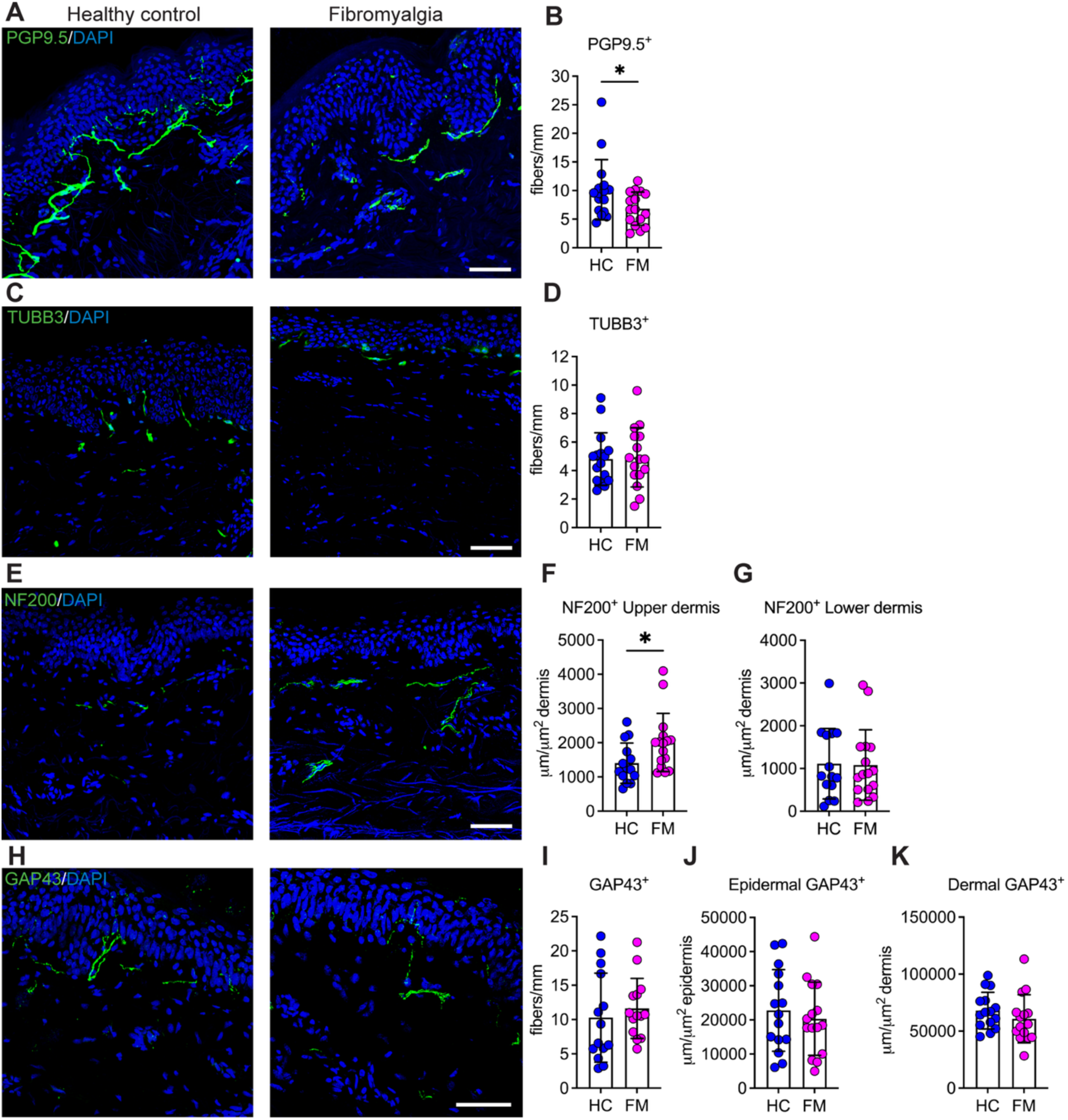
Characterization of epidermal and dermal profiles on FM skin. Skin biopsies from HC and FM were processed by immunohistochemistry against different markers. PGP9.5 was used to quantify the IENFD in HC and FM skin (A), where FM skin showed a reduction in the number of fibers/mm compared to HC (B). By using a cytoskeletal neuronal marker as TUBB3 (C) instead of a cytoplasmatic one, we observed no significant differences in the fibers/mm in the epidermis of FM patients compared to HC (D). The total length of nerve fibers expressing NF200 was analyzed in HC and FM skin (E). An increased length of NF200 fibers was found in the upper dermis (F), but a similar length was found in the lower dermis (G) in FM skin compared with HC. Finally, the skin was stained against GAP43 (H). The density of IENF positive for GAP43 was comparable between groups (I). Similarly, non-statistical differences were observed between HC and FM skin for the total axonal length of GAP43 fibers in the epidermis (J) and dermis (K). Data are shown as mean ± SD and analyzed by Mann-Whitney test, HC N=14, FM N=14-16, * p-value < 0.05. Scale bar = 50 μm.

The axons crossing the dermal-epidermal border are predominantly unmyelinated thin fibers. To also investigate potential changes in myelinated fibers, we assessed the total length of 200 kD neurofilament (NF200)-positive fibers in the skin of HC and FM subjects (Fig. 3E). The quantification showed that the total length of NF200^+^ fibers was significantly increased in the upper dermis of FM skin biopsies (2006 ± 849 µm/µm^2^) compared with HC (1403 ± 587 µm/µm^2^, p = 0.04) (Fig. 3F). However, no significant differences were observed in the lower dermis (Fig. 3G).

Growth-associated protein 43 (GAP43) is involved in axonal growth and regeneration (32, 33). Several studies have reported alterations in GAP43-immunoreactive cutaneous nerve fibers in patients with peripheral neuropathies(34, 35). Based on these findings, we examined the IENFD and the total length of axons expressing GAP43 in the epidermal and dermal regions (Fig. 3H). Our analysis showed that the density of GAP43^+^ fibers in the epidermis was similar in FM and HC (11.6 ± 4.4 and 10.3 ± 6.5 fibers/mm, respectively, p = 0.33, Fig. 3I). Similarly, the total GAP43^+^ axonal length in both the epidermis (Fig. 3J) and dermis (Fig. 3K) did not differ significantly between FM and HC skin biopsies.

### Reduced Schwann cells density in FM skin without changes in blood vessel length or area

A previous study reported a reduction in Schwann cells at the dermal-epidermal border— known as nociceptive Schwann cells—in skin samples from patients with small fiber neuropathy, a condition characterized by decreased distal IENFD (36). This finding prompted us to analyze the density of these cells in FM and HC skin. Nociceptive Schwann cells were identified as S100B-expressing cells that were negative for the Langerhans cell marker CD207 along or just below the dermal-epidermal border (Fig. 4A, C). Melanocytes expressing S100B were excluded by their characteristic morphology and location at the basal layer. Our analysis revealed no significant differences between FM and HC in the total density of S100B^+^CD207^-^ cells (Fig. 4B) or the ones associated with PGP9.5^+^ nerve fibers (presumably nociceptive Schwann cells) (Fig. 4D). However, we observed a lower density of non-nerve-associated Schwann cells along the dermal-epidermal border in the FM group (110.4 ± 48 cells/mm²) compared to HC (154.6 ± 47 cells/mm², p = 0.011; Fig. 4E).

**Figure 4.**
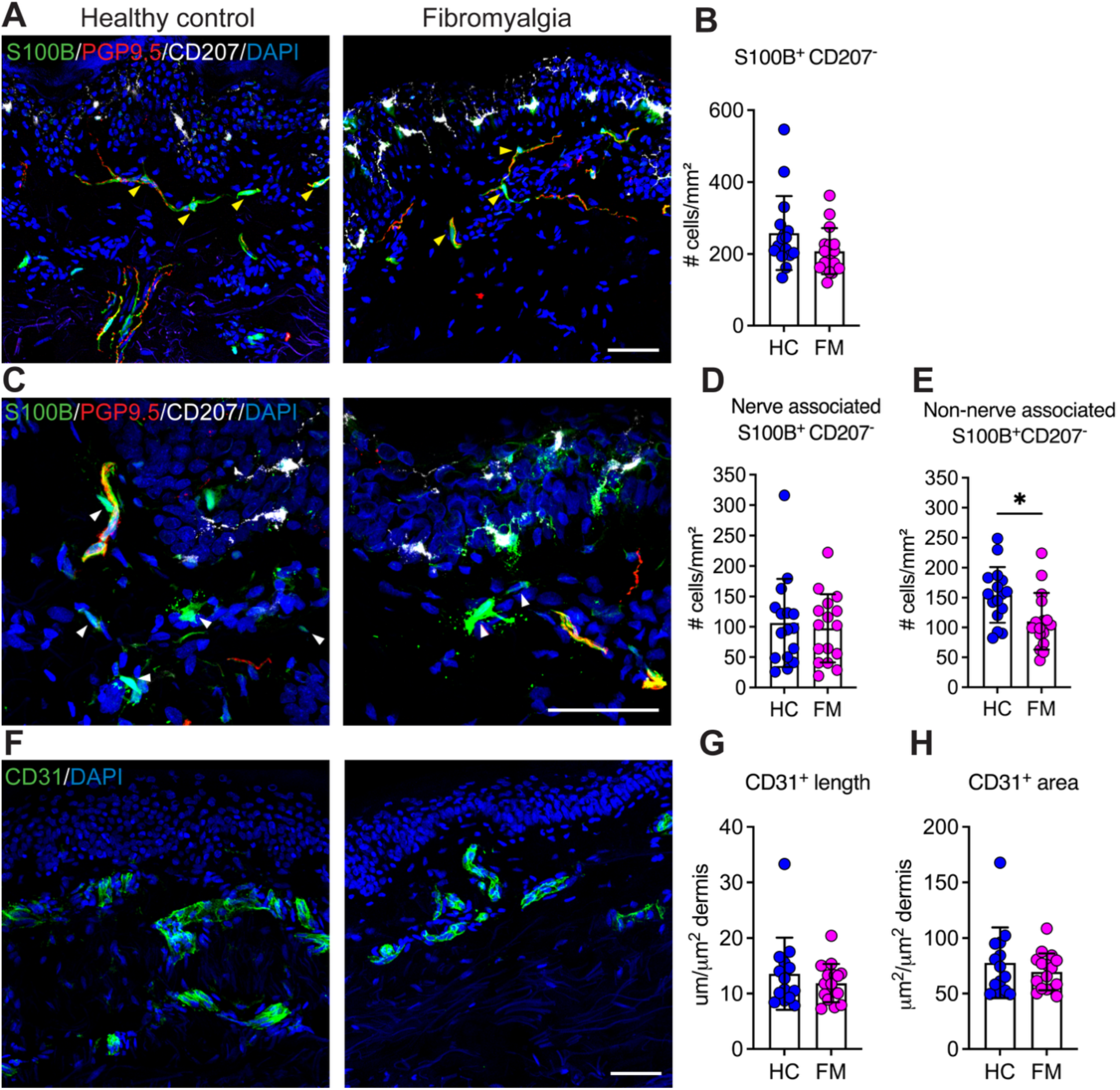
Quantification of Schwann cells and blood vessels in FM skin. S100b+ and CD207-cells were identified as dermal Schwann cells in skin biopsies from HC and FM volunteers (A). No changes in the S100b+CD207-total cell density were observed between groups (B). For further characterization, S100b+CD207-cells were subdivided to nerve-associated (yellow arrow heads) and non-nerve-associated (white arrow heads) by using PGP9.5 staining in skin biopsies from HC and FM participants (A, C). Nerve-associated S100b+CD207-cells (D) were similar between FM and HC. However, lower non-nerve associated S100b+CD207-cells were detected in FM skin (E). For analyzing length and density of blood vessels, we used CD31 (F). With this approach, we found that the length (G) and area (H) of CD31+ cells were similar between HC and FM groups. Data are shown as mean ± SD and analyzed by Mann-Whitney test, HC N=14, FM N=16, * p-value < 0.05. Scale bar = 50 μm.

A decrease in blood vessel density, in parallel with reduced IENFD, has been reported in individuals with FM (27). To determine whether similar vascular changes are present in our FM cohort, we labeled endothelial cells using an anti-CD31 antibody and assessed both the length and area of CD31^+^ vessels in the dermis (Fig. 4F). The results showed no significant differences in either the total length (Fig. 4G) or the area covered by CD31^+^ vessels (Fig. 4H) between FM and HC biopsies.

### FM skin shows no alterations in the density of Langerhans cells, dendritic cells or melanocytes

Langerhans cells, dendritic cells, and melanocytes are closely associated with sensory nerve fibers in the skin. The factors secreted by these cells can influence neuronal function, and changes in the secretion of neurotrophins and neurotrophic factors may regulate peripheral nerve fiber density in the epidermis and dermis (37–39). To investigate whether the density of these cells is altered in FM skin, we used antibodies targeting CD207 (langerin) for Langerhans cells (Fig. 5A), showing that the density of epidermal CD207^+^ cells was comparable between FM and HC samples (Fig. 5B). Additionally, Melan-A was used as a marker for melanocytes and CD1a for dendritic cells (Fig 5C). Melanocytes showed the characteristic uniform arrangement along the basal membrane, with no difference in melanocyte density between FM and HC biopsies (Fig. 5D). Lastly, there was no significant difference in the density of CD1a^+^ dendritic cells between the two groups, either in the epidermis or dermis (Fig 5E, F).

**Figure 5.**
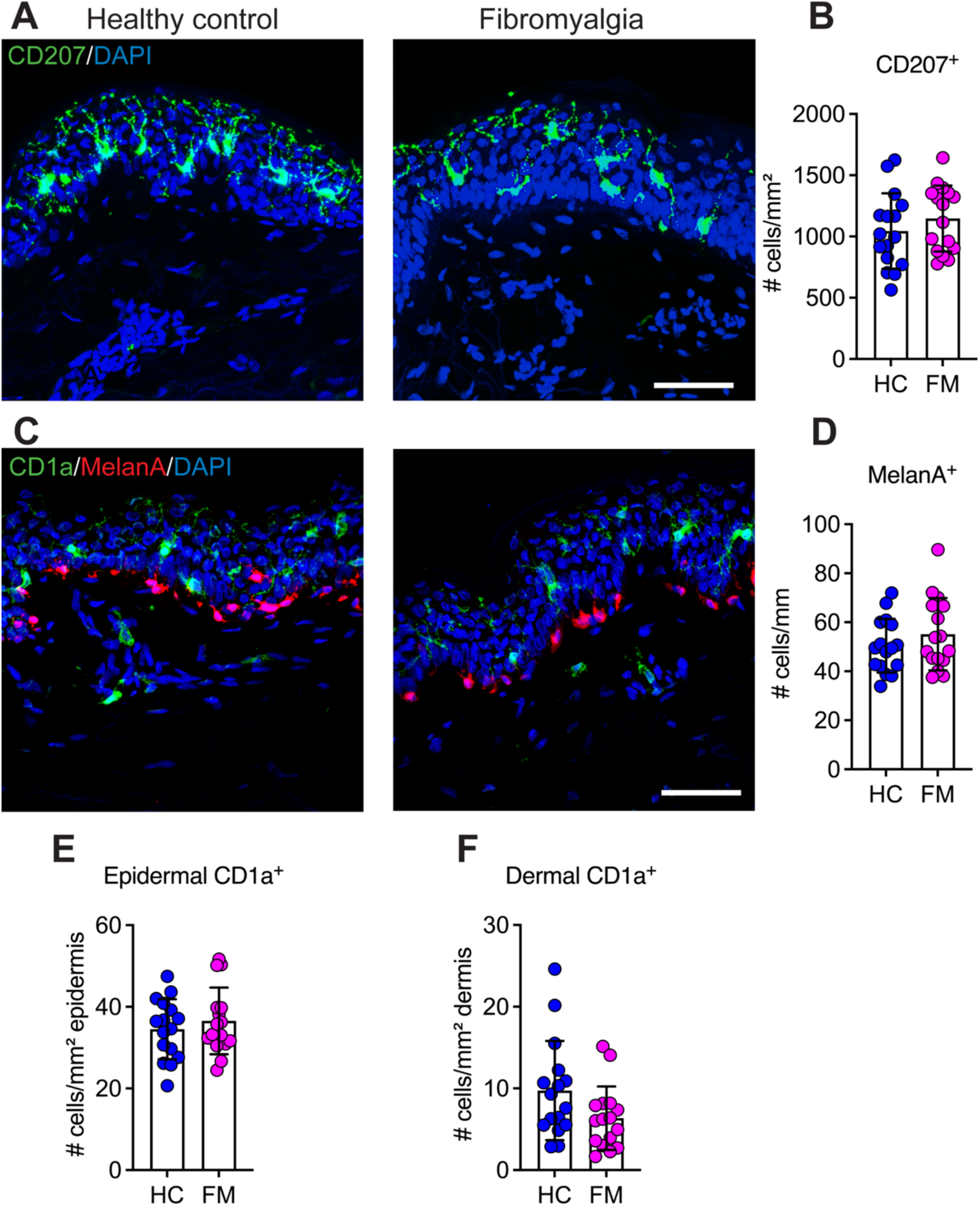
Density of dendritic cells in skin biopsies of FM and HC participants. Immunohistochemistry analysis of CD207+ cells in HC and FM skin (A) showed similar density of Langerhans cells between groups (B). To characterize the population of dendritic cells (DC), anti-CD1a and anti-MelanA antibodies were used to identify myeloid DC and melanocytes respectively, in biopsies from HC and FM skin (C). MelanA+ cells, corresponding to melanocytes in the dermal/epidermal border, showed no alterations in terms of density in FM skin compared to HC (D). Similarly, the density of CD1a+ cells was similar between groups in both the epidermis (E) and dermis (F). Data are shown as mean ± SD and analyzed by Mann-Whitney test, HC N=16, FM N=16, Scale bar = 50 μm.

### Dermal mast cell density is increased in FM skin without alterations in neutrophil number

In the dermis of individuals with FM, an increased number of mast cells has been previously reported (28), potentially contributing to skin sensitivity and low-grade inflammation. In our study, dermal mast cells were identified by the expression of FcɛRI and CD117 (cKit) (Fig. 6A). Our results showed that the number of FcERI^+^CD117^+^ was significantly increased in FM (154.3 ± 46.54 cells/mm^2^) compared to HC (114.1 ± 34.4 cells/mm^2^, p = 0.015) skin biopsies (Fig. 6B).

**Figure 6.**
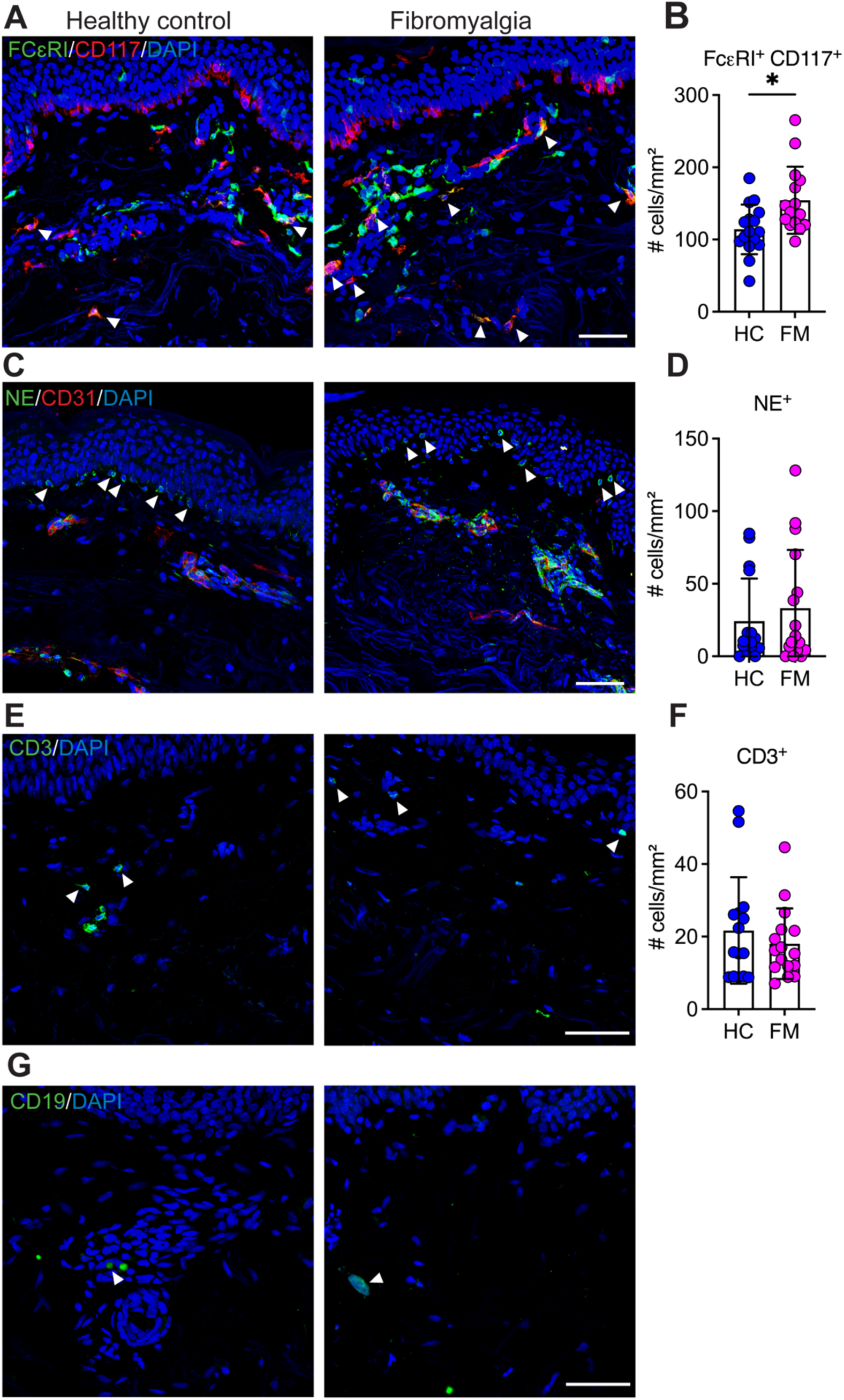
Quantification of different subsets of immune cells in FM skin. Double-positive FceRI+CD117+ cells (white arrows) were quantified in the dermis of HC and FM patients (A). The results showed a significant increase in mast cells in FM compared with HC biopsies (B). Neutrophils were identified by the expression of neutrophil elastase (NE) and overlap with a nuclear marker (DAPI), as well as the non-colocalization with endothelial cells (CD31) in HC and FM samples (C, white arrows). NE+ cells had a similar density between HC and FM patients (D). T cells were identified using an antibody against CD3 (E, white arrows). CD3+ cells were comparable in HC and FM skin (F). B cells were identified with a CD19 (G) but showed low density in both FM and HC samples, limiting meaningful quantitative analysis. Data are shown as mean ± SD and analyzed by Mann-Whitney test, HC N=15, FM N=15, * p-value < 0.05. Scale bar = 50 μm.

An increased number of mast cells in the skin is often associated with a prolonged inflammatory response, which may contribute to the recruitment of neutrophils. To determine whether this occurs in FM, we assessed neutrophil presence in extravascular tissue. This was done by analyzing the colocalization of neutrophil elastase (NE) with DAPI while ensuring no overlap with CD31, an endothelial marker, to differentiate circulating cells from those within the dermis and epidermis (Fig. 6C). Our analysis revealed a NE+ staining pattern on the dermal-epidermal border and epidermis, with no significant difference in the density of NE^+^ cells between FM patients and HC (Fig. 6D).

To investigate potential alterations in dermal adaptive immune cells in FM, we examined the presence of T cells and B cells. T cells were identified using an anti-CD3 antibody (Fig. 6E), and the quantification revealed no significant differences in dermal CD3^+^ cell density between FM and HC (Fig. 6F). B cells were detected using a CD19 antibody; however, B cell density was low in both FM and HC samples, with fewer than 2 cells per ROI and average (Fig. 6G), limiting the ability to perform meaningful quantitative comparisons.

### Dermal monocytes and macrophages are less abundant in FM skin

Although neutrophil numbers remained unchanged in skin biopsies, a recent study suggests they contribute to pain in FM (40). Given that neutrophils can influence macrophage polarization and that resident macrophages may play a critical role in neuronal homeostasis (41, 42), we next quantified the density of CD68^+^ cells in the dermis. Our analysis revealed a significant reduction of monocyte-derived cells in FM subjects (730 ± 209.4 cells/mm^2^) compared with HC (1045 ± 139 cells/mm^2^, p = 0.0002) Fig. 7A). Furthermore, we evaluated the morphology of these cells, including flatness and elongation (Fig. 7C). Dermal CD68^+^ cells in FM skin appeared slightly flatter compared to HC (Fig. 7D) and significantly more elongated (FM 2.3 ± 0.13 a.u vs HC 2.5 ± 0.2 a.u, p = 0.03) (Fig. 7F).

**Figure 7.**
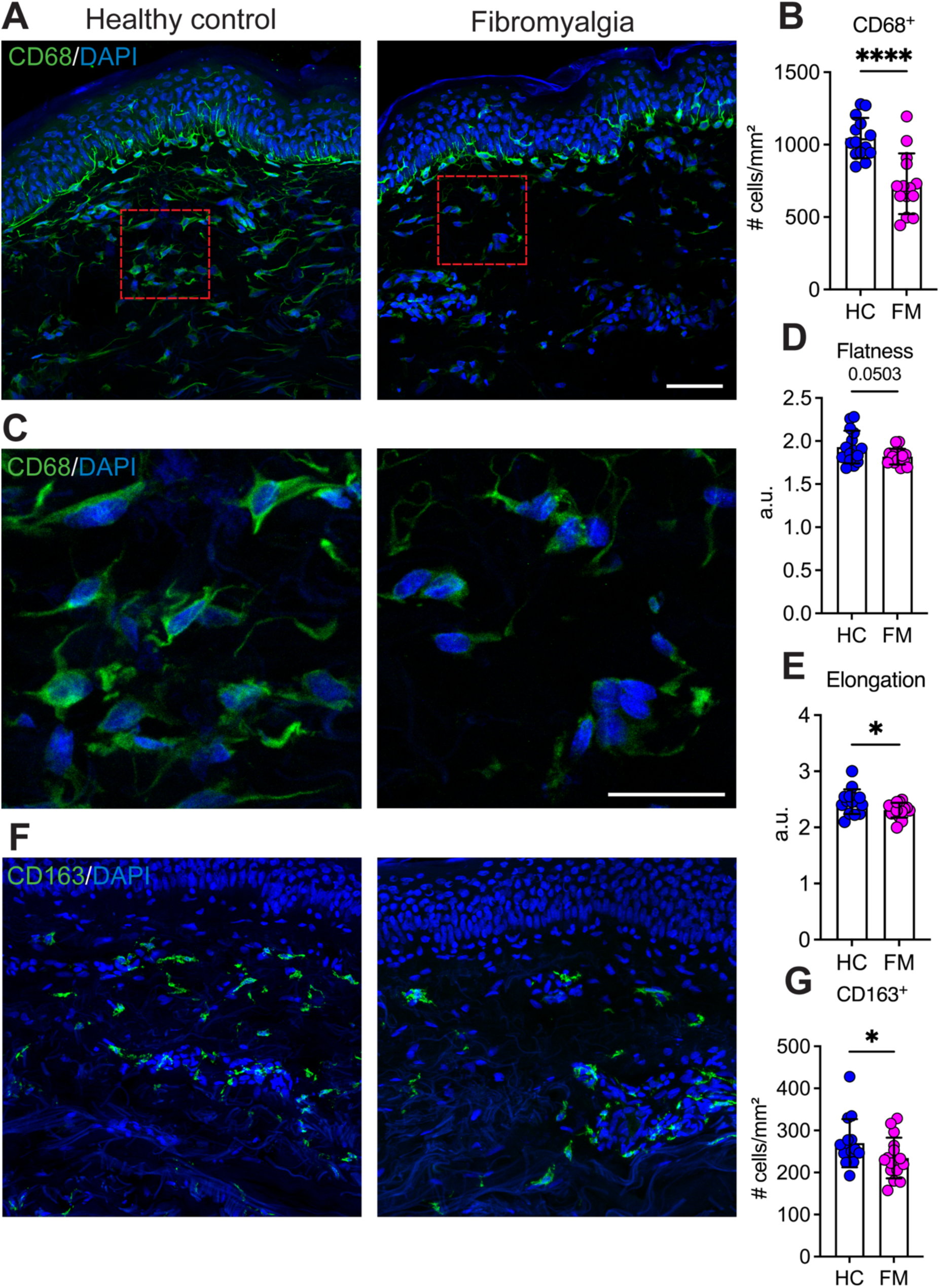
Density and morphology of macrophages in the dermis of FM biopsies. CD68+ cells were identified in the skin of HC and FM volunteers (A). A significantly lower density of CD68+ cells was found in the dermis of FM samples compared to HC (B). Changes in cell shape were also detected in FM CD68+ cells (C). While flatness was not statistically different (D), cells were less elongated in FM skin (F). To analyze a subset of macrophages, we quantified the density of CD163 positive cells (E). With this, a significantly lower density of CD163+ dermal macrophages was detected in FM skin compared with HC (G). Data are shown as mean ± SD and analyzed by Mann-Whitney test, HC N=14-15, FM N=14-16, * p-value < 0.05, **** p-value < 0.0001. Scale bar = 50 μm.

These findings raise the question of whether the reduction in CD68^+^ cells was related to a specific sub-population of macrophages. To explore this, we quantified CD163^+^ cells in skin samples, as this marker has been used to identify a subpopulation of tissue-resident macrophages in human skin and also, macrophages actively monitoring highly vascularized tissues (43–45) (Fig. 7E). Our analysis revealed a significant decrease in CD163^+^ cells density in the dermis of FM samples (234.4 ± 28.3 cell/mm^2^) compared to HC (269.7 ± 57.4 cells/mm^2^, p = 0.04) (Fig. 7G), underscoring a potential role of this subset in FM-associated pain hypersensitivity or disease severity.

### Fibromyalgia-specific associations between cutaneous cell populations and disease parameters

To test this hypothesis, we performed correlation analyses using the data from the thigh of FM patients and HC (Fig. 8). Pain thresholds related to skin sensitivity (HPT and CPT) were not significantly associated with PGP9.5+ IENFD in FM or HC, while PPT and VAS average pain were not significantly associated with any of the cell populations in both groups. We noted that the skin of HC showed a negative association between PGP9.5 IENFD and CD68+ cells (r = −0.56, p < 0.05), and a positive association with the total density of Schwann cells (r = 0.54, p < 0.05). Additionally, a strong negative association was seen between HPT and NF200 fibers in the lower dermis of FM skin (r = −0.75, p < 0.01), which was absent in HC (r = −0.46). For S100+ Schwann cells in FM skin, a negative association was seen against HPT (r = −0.53, p < 0.05), while this population was positively associated with CPT (r = 0.67, p < 0.01). This result was not significant in the skin of HC (Schwann cells vs HPT r = −0.13; Schwann cells vs CPT r = −0.31).

**Figure 8.**
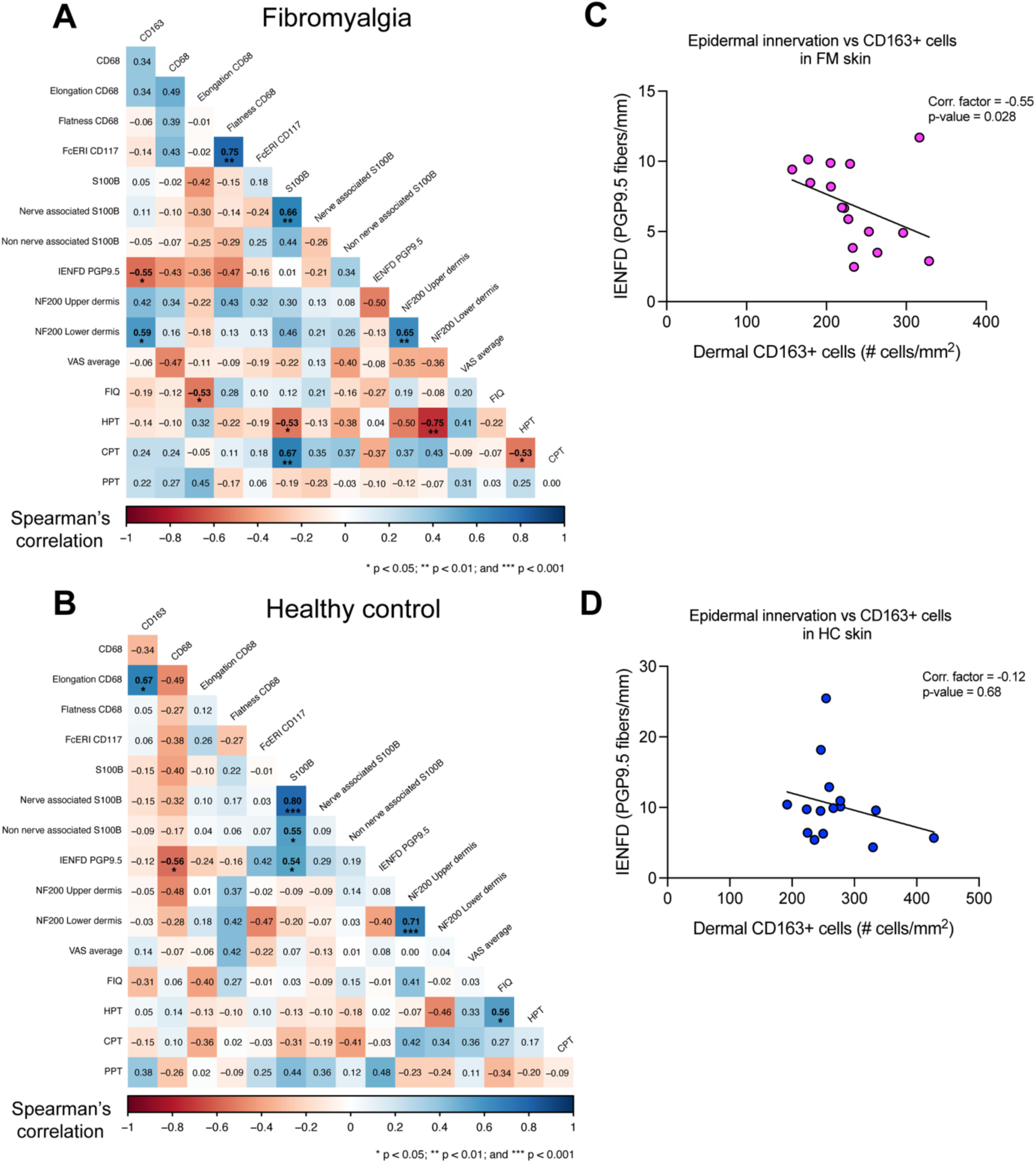
Correlation between cell populations, pain thresholds and disease severity. Using data from the thigh, correlation analyses were performed to examine the relationships between the most relevant findings in cutaneous cell populations and CPT, HPT, PPT, VAS pain levels, and FIQ in FM (A) and HC (B). The number and color in each square indicate the Spearman rank correlation coefficient, with statistically significant correlations (p < 0.05) shown in bold. Based on the negative correlation between PGP9.5+ IENFD and CD163+ cells in FM skin, we extended this exploratory analysis by visualizing the data distribution in FM (C) and HC (D), confirming a moderately strong association between epidermal innervation by PGP9.5+ fibers and dermal CD163+ macrophages in the skin of FM subjects. HC: N = 14– 15; FM: N = 14–16. Abbreviations: CPT, cold pain threshold; HPT, heat pain threshold; FIQ, Fibromyalgia Impact Questionnaire; PPT, pressure pain threshold.

Notably, FIQ scores were negatively associated against the elongation of CD68 cells (r = −0.53, p < 0.05), but not against flatness or other cell populations. The length of NF200 fibers in the upper and lower dermis were significantly and positively associated both in FM (r = 0.65, p < 0.01) as in HC skin (r = 0.71, p < 0.001). Finally, the density of dermal CD163+ cells was positively associated with the length of NF200+ in the lower dermis (r = 0.59, p < 0.05) in FM. Against PGP9.5+ IENFD, CD163+ cell were significantly and negatively associated in FM skin (r = −0.55, p < 0.05), but not in HC (r = −0.12; Figure 8C, D).

## DISCUSSION

Recent research highlights abnormalities such as heightened sensitivity and reduced epidermal innervation in FM, emphasizing the importance of investigating skin-related manifestations. Our study provides a comprehensive sensory and immunohistochemical characterization of FM skin compared to sex-matched HC. As expected, FM patients had a significantly increased multimodal pain sensitivity across different body sites (46). Histological examination revealed alterations in several cell populations within the epidermis and dermis. In the epidermis, FM skin showed a reduced density of PGP9.5⁺ small fibers. Surprisingly, the dermis displayed notable changes; first, increased NF200+ fiber density in the upper dermis of FM skin, a decrease in non-nerve associated Schwann cells, CD68+ and CD163+ macrophages, as well as an increase in mast cell count. These findings highlight distinct cellular alterations in FM skin, particularly within the dermal compartment.

In addition to reduced pressure pain thresholds, QST revealed abnormalities in heat and cold pain thresholds, indicating thermal and mechanical hyperalgesia at two different locations. Importantly, detection of warm and cold temperatures remained intact in this cohort of FM patients at both the arm and the thigh. FM patients often show signs of small fiber pathology in a non-length-dependent manner (8, 47, 48), which we found here as the biopsies were collected from the upper hip area. While our findings showed preserved thermal detection—consistent with previous studies (49, 50)—other reports have described altered warm, cold, and mechanical detection thresholds in FM patients (8). These discrepancies may reflect distinct sensory profiles among FM patients with and without hypoesthesia (51).

Even though pain in FM is typically associated with diffuse, deep pain sensations (52), few studies have noted burning sensations as prevalent in FM (13, 51, 53). Here, over 80% of the FM subjects reported burning and stinging, broadening the clinical profile and emphasizing that FM includes a spectrum of pain qualities beyond deep pain and sensitivity to mechanical and thermal stimuli.

A reduction in intraepidermal innervation in chronic pain conditions, including FM, has been extensively reported, mostly using PGP9.5 as marker (8–11, 54). Our results align with these prior reports and reinforce the concept of small fiber pathology in FM (48, 51). To further characterize skin innervation, we also used the cytoskeleton marker TUBB3 to quantify the density of IENF. However, we did not find a difference between FM and HC samples. We noted that TUBB3 is expressed in the soma and dendritic processes of other cells in the epidermal basal layer, such as melanocytes (55), which complicates the clear identification of crossing axons. These limitations support the use of PGP9.5 in counting IENF and may explain discrepancies observed between the two markers.

Most myelinated sensory fibers, including both nociceptive and non-nociceptive, express NF200 (56). Notably, there was an increased length of NF200+ nerve profiles in the upper dermis in FM biopsies compared to controls. A previous report did not detect differences in the NF200+ fibers in FM dermis (57), likely due to differences in biopsy site and assessment method. In our study, skin samples were collected from the upper thigh, whereas earlier research examined the lower calf. Increased innervation in the palmar glabrous skin of FM patients has been reported, specifically around arteriole-venule shunts (58). Although vascular innervation was not our focus, we quantified the length and volume of CD31+ vessels, finding no differences between FM and HC. While this aligns with other reports indicating no change in blood flow or vascular reactivity in FM skin (59), some studies showed reduced peripheral blood flow and altered capillary dilatation in FM patients (60).

GAP43 (neuromodulin) is a highly expressed protein on growth cones during nerve regeneration and sprouting (33). In our study, epidermal and dermal axonal lengths and crossings of GAP43-expressing fibers were similar between FM subjects and HC. This contrasts with a previous report showing lower IENFD of GAP-43 positive fibers in the lower leg and proximal thigh of FM subjects (8), likely due to differences in thickness of cryostat sections.

Skin innervation is regulated by a fine balance of guidance cues from resident cells (21, 57, 61). For example, increased expression of ephrin-A4 (*EFNA4)* and its receptor ephrin-receptor-A4 (*EPHA4)* was found in fibroblasts and keratinocytes from FM patients (27). These molecules guide axon growth, EFNA4 attracts, and EPHA4 repels axons (61, 62), potentially contributing to the distinct innervation patterns observed in FM skin. Although we did not assess these types of cues, our findings highlight the need for further research into nerve guidance signals and growth factors across skin layers in FM.

Recent findings have identified subepidermal Schwann cells that initiate nociceptive responses to mechanical stimuli in mice (63), and their presence in human dermis has now been confirmed (23). These cells are characterized by S100B expression, lack of CD207 expression that excludes Langerhans cells, and proximity to neurons near the epidermis. In our study, we quantified S100B+ cells in the dermis and using PGP9.5 as a pan-neuronal marker, we also distinguished nerve-associated from non-nerve associated Schwann cells. Unlike reports in small fiber neuropathy (36), we found no reduction of nerve-associated S100B+CD207-cells in FM skin, but a lower density of non-nerve associated S100B+ cells compared to HC. Schwann cells have been implicated in tissue remodeling and wound healing (64, 65), especially during inflammation, proliferation, and extracellular matrix (ECM) remodeling (45). Abundant in hypertrophic scars (keloids), non-nerve Schwann cells often display de-differentiation and pro-fibrotic characteristics, promoting ECM overproduction (65). Therefore, the reduced non-nerve associated S100B+ cells in FM skin may indicate impaired tissue support and repair during critical ECM-dependent healing phases (45, 65).

In our study, the density of dendritic cells (CD1a+ and CD207+) was similar between FM and HC, indicating no major alterations in epidermal immune surveillance (38). Recently, it was described that the density of Langerhans cells is increased in patients with painful diabetic neuropathy (25). Given no changes were seen in our cohort of FM, this suggests a potential key difference between cell-related pain mechanisms related to neuropathic vs nociplastic pain. Density of melanocytes was also similar between FM patients and controls, indicating that, despite proposed associations between FM and vitiligo (18, 66), key epidermal immune cell populations remain unaffected in FM skin.

Although limited, some studies report pruritus as a symptom in FM (13, 53). As a common dermatological symptom, pruritus is closely linked to mast cells, which express FceRI and synthesize histamine, a potent pruritogen upon release (67). Co-expression of c-Kit (CD117) further characterizes skin mast cells (68). We observed an increased number of FceRI/CD117 positive mast cells in dermis, aligning with previous reports of mastocytosis in FM skin (28, 69). However, a phase 1 randomized controlled clinical trial with ketotifen, a histamine H1 receptor antagonist and mast cell stabilizer, did not improve pain intensity in FM (70). These observations suggest that mast cells may contribute to overlooked symptoms like itch or burning sensations, potentially linking these sensory abnormalities to underlying immunological or neuroinflammatory processes.

Emerging research highlights circulating neutrophils as potential contributors to FM pathology. Their ability to infiltrate tissues in various conditions—such as autoimmune diseases, infections, and cancer (71–73)—and to influence pain sensitivity suggests a broader role for immune cells in modulating FM symptoms. For example, a recent study showed that transferring neutrophils from FM patients into mice induces widespread pain, with these cells observed in the DRG, indicating a possible contribution to FM (40). However, whether these cells were within blood vessels or DRG parenchyma was not addressed.

Neutrophil elastase (NE), a protease released upon neutrophil activation during inflammatory responses, is commonly used as a marker for neutrophils (71). In healthy skin, neutrophils are typically infrequent (71, 74). However, a recent study identified a specific population of collagen-producing neutrophils enriched in human skin (75). In autoimmune diseases such as systemic lupus erythematosus, neutrophils are frequently found around blood vessels, hair follicles, and eccrine glands (74). They can also be present in the papillary dermis, in close contact with the dermal–epidermal junction (74). To ensure accurate quantification of neutrophils in the skin, we included CD31 staining to exclude NE⁺ cells colocalized with this vascular marker. In our cohort, we found no significant differences in NE⁺ cell density in FM skin compared to controls. However, it is important to note that NE⁺ cells were predominantly located along the basal membrane and even within the epidermis, suggesting that not all NE⁺ signals necessarily correspond to neutrophils. Given neutrophils’ short lifespan and the absence of clear skin inflammation in FM, their impact may be more relevant in regions like the DRGs, where they could directly influence pain mechanisms.

Notably, increased expression of NK cell activation ligands has been observed in skin biopsies from FM patients, along with the presence of NK cells near peripheral nerves (30), suggesting a role of these cells in pain induction. We did not assess NK cells in the current study but focused on CD3+ T cells as studies of T cells in circulation in FM suggest altered frequency and polarization, primarily within the CD4+ subset, though findings are inconsistent (76). Our analysis showed no difference in CD3+ cell density in the skin between FM patients and HC. B cells are present in healthy skin but typically at low densities. In our study, we observed occasional CD19+ B cells, with overall numbers being very low and not different between groups when assessed qualitatively.

Increased nerve remodeling and skin healing processes can promote the activation and recruitment of phagocytic cells (45, 77). Importantly, alterations in the local cutaneous cellular landscape, whether in cell number or function, may contribute to various pathological processes (41, 78, 79). For example, higher density of macrophages/monocytes are typically associated with lesional skin in conditions like atopic dermatitis, discoid lupus erythematosus, and psoriasis (80–82), but also with keloid scarring (65, 83). Contrarily, we observed a reduction in the number of dermal CD68+ cells in FM skin. Moreover, the subset of macrophages expressing the scavenger receptor CD163 was significantly decreased. Interestingly, recent studies have shown that thymic CD68+ and CD163+ macrophages were significantly decreased in Myasthenia gravis patients (84), a chronic autoimmune disorder that results in muscle weakness. The mechanisms and implications of these decreases remain elusive. In animal models, depletion of dermal macrophages impairs wound healing (85), while in human skin and organotypic models, gene expression of CD68 and CD163 increases during wound healing (45) and seems to promote more efficient re-epithelialization, respectively (79).

Additionally, we found that CD68+ cells presented morphological changes in FM skin compared to HC, appearing flattened and less elongated. Cell shape is relevant for modulating polarization and motility of macrophages (86, 87). For instance, in vitro treatment with LPS and IFN-γ induces morphological changes toward a more rounded shape in bone marrow–derived macrophages (86). In contrast, elongated cell shapes are associated with enhanced tissue healing programs (83, 86, 87). Overall, the reduced number of CD68⁺ and CD163⁺ cells in FM skin, plus the morphological alterations, may therefore indicate a compromised cellular response, impaired tissue repair processes or resolution of inflammation (83). Moreover, our group recently identified a specialized role for CD163+ macrophages in surveying vascular content at the blood-DRG barrier, with altered responses during systemic inflammation (43). Whether the lower number of CD163+ expressing cells in FM skin hampers wound healing skin barrier function warrants further investigation.

As a hypothesis-driven explorative analysis, we performed correlation analyses between the most relevant cell populations of this study and thermal pain thresholds as measures of pain sensitivity, PPT for mechanical hypersensitivity, and FIQ scores and average VAS average pain levels as variables related to disease severity. In our cohort, PGP9.5 IENFD did not correlate with QST parameters, FIQ scores, or pain levels, consistent with reports from other pain conditions where nerve fiber loss does not translate to higher pain sensitivity (88, 89). Within FM skin, we also noted that the length of NF200 fibers in the lower dermis correlated negatively with HPT. Even though our results showed increased length patterns of NF200+ fibers in the upper dermis of FM patients, since this association was absent in HC, a link between Aδ fibers and heat pain thresholds in FM is possible.

Schwann cells are linked to sensing noxious stimuli in the skin (23, 63). In our analysis, the density of Schwann cells correlated negatively with HPT and positively with CPT, suggesting a broader nociceptive involvement in response to thermal stimuli beyond mechanosensation in FM.

Previous reports have shown that metabolic and mitochondrial dysfunction can cause small fiber injury and macrophage loss (90, 91). Furthermore, in animal models of diabetic peripheral neuropathy, recruitment of macrophages results in a neuroprotective effect that delayed loss of sensory axons in the sciatic nerve. Conversely, an accelerated skin denervation was observed with the inhibition of macrophages recruitment (92). In our correlation analyses, FM patients with lower IENFD tended to have higher CD163⁺ macrophage counts. Even though this result does not prove causality or a direct interaction between these cell population, it points to a relevant neuroimmune interaction between dermis and epidermis that can potentially modulate the skin innervation in FM.

Notably, we noted a negative association between FIQ scores and an elongated shape in CD68+ macrophages in FM. Since reparative macrophages switched to an elongated shape during healing (86, 93), this finding gives additional evidence that specific subsets of skin cell populations, particularly dermal macrophages, may have a functional alteration in the skin of FM patients, potentially impacting cutaneous repair processes (86, 87). Although exploratory and limited by sample size, this integration of cutaneous cells populations, sensory data, and clinical information from the same individuals is a rare and valuable resource for future larger studies.

One limitation of our study is the age difference between the FM and HC groups. However, this difference corresponds to approximately one decade, with the FM group having a younger age. Considering that the number of small fibers, melanocytes, and Langerhans cells, among other skin cell populations, normally decline with age, this may potentially lead to an underestimation of the cutaneous changes reported here (94–96). In this sense, it would be relevant in the future to study the phenotypic differences in younger and older cohorts of FM subjects, allowing us to further characterize the temporal dynamics of changes in the skin cellular landscape. Our focus on cell numbers provides valuable insights, but it is essential to recognize that functional changes and interactions, such as signaling pathways and immune responses, may occur without detectable differences in cell counts. These dynamic processes could play a significant role in FM, underscoring the need for further research to elucidate these functional aspects and assess their implications.

In summary, although skin-related symptoms have received less attention in FM research, likely due to the prominence of fatigue and widespread pain, our findings highlight the importance of considering cutaneous alterations, particularly within the dermis, as part of a broader understanding of FM. Furthermore, the complex interplay between nerve fibers and immune cell populations observed within the different skin microenvironments, contributes to the evidence of immune system involvement in FM pathology. However, no clear signs of inflammation were observed in FM skin. Finally, dissecting the role and the functional implications for skin homeostasis of our novel findings, namely, higher length NF200+ fibers in the upper dermis, reduced non-nerve associated dermal Schwann cells, and decreased density of CD68+ and CD163+ cells in the dermis, may have broader implications for tissue repair, pain mechanism, and potential treatments for subjects with FM syndrome.

## METHODS

### Participants

Given FM is significantly more prevalent in women than in men, this study group included 16 female subjects with FM and 16 healthy females who served as healthy controls (HC) (see Table 1). The subjects were recruited from a larger trial including imaging (NCT05815381). To ensure adherence with ACR1990 and ACR2016 classification criteria (2, 3) all FM subjects underwent systematic screening by a specialist in rehabilitation medicine. Inclusion criteria for FM included being right-handed, and of age 20–65 years. Exclusion criteria were: other dominant pain conditions than FM, other autoimmune or rheumatoid diseases, other severe somatic diseases (cardiovascular, neurological, diabetes mellitus, cancer, etc.), substance abuse, psychiatric disorders including ongoing treatment for anxiety or depression, medications with antidepressants, anticonvulsants or corticosteroids, previous heart or brain surgery, hypertension (> 160/90 mmHg), obesity (body mass index > 35), smoking (> 5 cigarettes/day), pregnancy, self-reported claustrophobia, magnetic implants, inability to speak and understand Swedish, being unable to refrain from hypnotics, analgesics, or non-steroidal anti-inflammatory drugs 48 hours prior to participating in the study, and contraindications for skin biopsy, *i.e.* hypersensitivity to local anesthetics, and hemophilia. HC were right-handed females, without regular medications with non-steroidal anti-inflammatory drugs, sleep medication or analgesics, as well as free from chronic pain and from the exclusion criteria stated above for FM. All participants were recruited through daily press advertisement. They were compensated for their time and participation.

### Clinical, demographic, and sensory characteristics

We collected demographic data on age, height, and weight; race and ethnicity were not collected. All participants reported their current (during the visit) and average pain levels during the last week using a 0-100 mm visual analogue scale (VAS now and VAS average, respectively), where 0 indicated no pain and 100, the worst possible pain. Additionally, subjects also reported the presence of pain descriptors, such as burning and stinging using the short form McGill pain questionnaire on a scale from 0 to 3 (none, mild, moderate, severe). Participants answered the Fibromyalgia impact questionnaire (FIQ), a disease-specific self-reported questionnaire that includes 19 items of disabilities and symptoms related to FM ranging from 0 to 100, with higher score indicates a lower quality health status(97).

### Quantitative Sensory Testing

Sensory thresholds were assessed on the dorsal side of the left arm and lateral side of the left thigh of each subject for the following parameters: warm detection threshold (WDT), heat pain threshold (HPT), cold detection threshold (CDT), and cold pain threshold (CPT). The method of limits was used with a Thermotest device (Somedic Sales AB, Sweden; 50 × 25 mm probe). For each anatomical site, three trials of continuously ascending temperatures were used for warm stimulation, and three trials of continuously descending temperatures were used for cold stimulation. The baseline temperature was 32°C, with a linear change in temperature ranging from 4-50°C, heating/cooling 1°C/s, back 5°C/s. A randomized interstimulus interval (4-6s) was used for the innocuous trials (WDT, CDT), while the noxious trials were started manually, with at least 30s intervals. Subjects were instructed to press a button as soon as they perceived the target sensation, either the slightest detection of temperature or the first sensation of pain.

Sensitivity to suprathreshold stimuli was determined for heat, cold, and pressure pain at the left forearm and thigh using a previously described protocol with one assessment per site and intensity (98). Briefly, subjects were asked to indicate when the temperature or pressure reached an intensity rated as 4/10 (moderate-to-strong pain) and 7/10 (very strong pain), respectively, on the Borg category ratio 10 scale (Borg CR 10) (99, 100).

Pressure pain threshold (PPTs) were determined using a hand-held algometer (Somedic Sales AB, Sweden) with a rubber probe of 1 cm^2^, and with an approximate rate of pressure increase of 50kPa/s following our previously reported protocol (101). Subjects were first familiarized with the procedure and instructed to press a response button at their first sensation of pain. PPTs were assessed bilaterally at the supraspinatus, and the gluteus muscle, once per site and the arithmetic mean was calculated across sites. In addition, PPTs were assessed three times at the left forearm and the left thigh, respectively.

### Skin biopsies

Skin biopsies (4-mm punch) were collected from each subject with aseptic technique after local anesthesia from the lateral side of the upper thigh. The biopsies were immediately frozen in dry ice and stored at −80 °C until processing.

### Immunohistochemistry

Skin samples were incubated in 2 ml of a pre-cooled 4% formaldehyde solution at 4°C for 2 hours for tissue fixation. Then, samples were transferred to a 30% sucrose solution for cryoprotection. Once the skin sank in the tube, individual OCT blocks were prepared and stored at −20°C until cryocutting. Twenty-micron sections were collected in superfrost plus slides and stored at −20°C until processing. For immunohistochemistry (IHC) assays, the skin sections were let dry at room temperature for 30 minutes and re-hydrated in 0.1M PBS for 10 minutes, followed by 0.01M PBS for 10 minutes. Then, the sections were blocked with a solution containing 3% normal donkey serum (NDS, S30-M, Merck, Germany) and 0.3% triton X-100 (X100, Sigma, Germany) in 0.01M PBS for 1 hour. Next, the slides were incubated overnight with antibodies against CD1a, CD3, CD19, CD31,CD68, CD163, CD207, GAP43, MelanA, NE, NF200, PGP9.5, S100B, and TUBB3 in different cocktail combinations (details in Supplementary Table 1). On the second day, the slides were washed three times with 0.01 M PBS and incubated with the appropriate secondary antibodies conjugated with Cy2, Cy3, Cy5, or Alexa Fluor 488 and counterstained with 4’,6-Diamidino-2-Phenylindole, Dihydrochloride (DAPI, 1:20000, D1306, Invitrogen, United Kingdom). Finally, the slides were dehydrated in an alcohol gradient, cleared with xylene twice, and coverslipped with permanent mounting media (Pertex, 00811, Histolab, Sweden). Additionally, another set of slides was incubated overnight with FCeRI-PE and CD117-APC, followed by counterstaining with DAPI and coverslipped with ProLong™ Gold Antifade Mountant (P36934, Invitrogen, Sweden).

### Image acquisition and analysis

Images were acquired using a Zeiss LSM800 confocal system with Zen 3.6 software. A random ROI in the biopsy was acquired using a 20X objective. Maximum projections of z-stacks were acquired and used for IENFD and immune cell densities analyses, and the average of 4 consecutive sections was calculated. IENFD was assessed following the EFNS guidelines on the use of skin biopsy in the diagnosis of peripheral neuropathy(54). Individual nerve profile crossings were counted, and the total number was divided by the basement membrane (BM) length. For immune cells, DAPI was used to identify individual cells, and the total number of cells was divided by the epidermal or dermal area. The experimenters were blinded to the subject condition. Gap43, NF200, blood vessel (BV) length, macrophage shape, analyses were done using the drgquant pipeline (102). Briefly, U-NET (103) models were trained to identify vasculature, macrophages, axons, and identify the epidermis, dermis, and stratum corneum. Model outputs were then run through macros in FIJI (104) that segmented structures using connected component analysis in CLIJ (105). Data tables were then concatenated using scripts written in Python. For NF200 and BV, the whole biopsy was imaged, and the average of 4 different sections was calculated. For NF200 axonal length, the dermis was divided into upper dermis (150 μm from the BM) and deep dermis (more than 150 μm).

### Statistical analysis

All data is presented as the mean and standard deviation (SD). According to the data statistical distribution, a Mann-Whitney U test was used to compare the two groups. For threshold and suprathreshold stimuli, two-way repeated measures ANOVA was used to analyze changes in groups and levels of pain on each volunteer. Spearman rank correlations were used to explore relationships between cell populations and clinical data. All data was graphed and analyzed using GraphPad Prism 10 (Boston, USA) except for correlations that were analyzed in R Studio version 2024.4.2.764.

### Study approval

This study was approved by the Swedish ethical committee (Etikprövningsmyndigheten, 2019-06161). Written informed consent was received from all subjects prior to participation, and the study was performed in accordance with the declaration of Helsinki.

### Data availability

Underlying data for figures and reported means are available in the Supporting Data Values file.

## Supporting information

Supplementary table 1. Antibodies and conditions used.

## AUTHOR CONTRIBUTIONS

Design of research study: CMU, EK, CIS

Recruitment and obtention of QST and samples: KS, EL, KE, SF, ML, EK, TA

Tissue processing: CMU, AK, SVL

Acquiring data: CMU, MH, KA, AK, SVL, MTB, KE, SF

Analyzing data: CMU, MH, MTB, KA, AK, SVL, EK, CIS

Writing the manuscript: MTB, CMU, EK, CIS

## ACKNOWLEDGMENTS

The authors thank Juan A. Vazquez-Mora for technical assistance. Swedish Research Council grant 542-2013-8373 (to C. I. Svensson) and grant 2022-00564 (to E. Kosek), the Knut and Alice Wallenberg Foundation (018.0161 to C. I. Svensson), the Karolinska Institute Foundations (to C. E. Morado-Urbina), the European Union Seventh Framework Programme (FP7/2007–2013) under grant agreement number 602919 (to C. I. Svensson and E. Kosek), European Union’s Horizon 2020 research and innovation programme under the Marie Skłodowska-Curie Grant Agreement number 764860 (to C. I. Svensson and E. Kosek), the European Research Council (ERC) under the European Union’s Horizon 2020 research and innovation programme under the grant agreement no. 866075 (to C. I. Svensson), FOREUM (to C. I. Svensson and E. Kosek), the Swedish Rheumatism Association (to E. Kosek and C.I. Svensson), by a generous donation from Leif Lundblad and family (to C. I. Svensson and E. Kosek), Stockholm County Council (grant ALF-20190039 and RS2023-0859 to E. Kosek) and a grant from Fibromyalgiförbundet (E.Kosek).

## REFERENCES

1. Sarzi-Puttini P, et al. Fibromyalgia: an update on clinical characteristics, aetiopathogenesis and treatment. Nat Rev Rheumatol. 2020;16(11):645–660.

2. Wolfe F, et al. The American College of Rheumatology 1990 Criteria for the Classification of Fibromyalgia. Report of the Multicenter Criteria Committee. Arthritis Rheum. 1990;33(2):160–72.

3. Wolfe F, et al. 2016 Revisions to the 2010/2011 fibromyalgia diagnostic criteria. Semin Arthritis Rheum. 2016;46(3). 10.1016/j.semarthrit.2016.08.012.

4. Kravitz HM, Katz RS. Fibrofog and fibromyalgia: a narrative review and implications for clinical practice. Rheumatol Int. 2015;35(7):1115–25.

5. Kosek E. The concept of nociplastic pain-where to from here? Pain. 2024;165(11S):S50–S57.

6. Pinto AM, et al. Neurophysiological and psychosocial mechanisms of fibromyalgia: A comprehensive review and call for an integrative model. Neurosci Biobehav Rev. 2023;151:105235.

7. Mawla I, et al. Large-scale momentary brain co-activation patterns are associated with hyperalgesia and mediate focal neurochemistry and cross-network functional connectivity in fibromyalgia. Pain. 2023;164(12):2737–2748.

8. Üçeyler N, et al. Small fibre pathology in patients with fibromyalgia syndrome. Brain. 2013;136(Pt 6):1857–67.

9. Caro XJ, Winter EF. Evidence of abnormal epidermal nerve fiber density in fibromyalgia: clinical and immunologic implications. Arthritis Rheumatol. 2014;66(7):1945–54.

10. Giannoccaro MP, et al. Small nerve fiber involvement in patients referred for fibromyalgia. Muscle Nerve. 2014;49(5):757–9.

11. Teng H-W, et al. Altered sensory nerve excitability in fibromyalgia. J Formos Med Assoc. 2021;120(8):1611–1619.

12. Serra J, et al. Hyperexcitable C nociceptors in fibromyalgia. Ann Neurol. 2014;75(2):196–208.

13. Laniosz V, Wetter DA, Godar DA. Dermatologic manifestations of fibromyalgia. Clin Rheumatol. 2014;33(7):1009–1013.

14. Kaya Erdogan H, et al. Cutaneous findings in fibromyalgia syndrome and their effect on quality of life. Dermatologica Sinica. 2016;34(3):131–134.

15. D’Onghia M, et al. Fibromyalgia and Skin Disorders: A Systematic Review. J Clin Med. 2024;13(15):4404.

16. Kulakli S, et al. Relationship Between Chronic Spontaneous Urticaria and Fibromyalgia Syndrome. Medical Records. 2023;5(3):638–43.

17. Yazmalar L, et al. High Frequency of Fibromyalgia in Patients With Acne Vulgaris. Arch Rheumatol. 2016;31(2):170–175.

18. Aşkın A, et al. Could There Be a Possible Link Between Vitiligo and Fibromyalgia Syndrome? Turkish Journal of Dermatology / Türk Dermatoloji Dergisi. 2018;12(2):100–106.

19. Ji R-R, Chamessian A, Zhang Y-Q. Pain regulation by non-neuronal cells and inflammation. Science. 2016;354(6312):572–577.

20. Pinho-Ribeiro FA, Verri WA, Chiu IM. Nociceptor Sensory Neuron-Immune Interactions in Pain and Inflammation. Trends Immunol. 2017;38(1):5–19.

21. Wang F, Julien DP, Sagasti A. Journey to the skin: Somatosensory peripheral axon guidance and morphogenesis. Cell Adh Migr. 2013;7(4):388–94.

22. Nguyen A V, Soulika AM. The Dynamics of the Skin’s Immune System. Int J Mol Sci. 2019;20(8). 10.3390/ijms20081811.

23. Rinwa P, et al. Demise of nociceptive Schwann cells causes nerve retraction and pain hyperalgesia. Pain. 2021;162(6):1816–1827.

24. Nestle FO, et al. Skin immune sentinels in health and disease. Nat Rev Immunol. 2009;9(10):679–91.

25. Pacifico P, et al. Epidermal Langerhans Cells Drive Painful Diabetic Neuropathy in a Sex-dependent Manner [preprint]. 2025. 10.1101/2025.01.30.635734.

26. Kosmidis ML, et al. Reduction of Intraepidermal Nerve Fiber Density (IENFD) in the skin biopsies of patients with fibromyalgia: a controlled study. J Neurol Sci. 2014;347(1–2):143–7.

27. Evdokimov D, et al. Characterization of dermal skin innervation in fibromyalgia syndrome. PLoS One. 2020;15(1):e0227674.

28. Blanco I, et al. Abnormal overexpression of mastocytes in skin biopsies of fibromyalgia patients. Clin Rheumatol. 2010;29(12):1403–12.

29. Conti P, et al. Impact of mast cells in fibromyalgia and low-grade chronic inflammation: Can IL-37 play a role? Dermatol Ther. 2020;33(1):e13191.

30. Verma V, et al. Unbiased immune profiling reveals a natural killer cell-peripheral nerve axis in fibromyalgia. Pain. 2022;163(7):e821–e836.

31. Sánchez-Domínguez B, et al. Oxidative stress, mitochondrial dysfunction and, inflammation common events in skin of patients with Fibromyalgia. Mitochondrion. 2015;21:69–75.

32. Ragé M, et al. The time course of CO2 laser-evoked responses and of skin nerve fibre markers after topical capsaicin in human volunteers. Clinical Neurophysiology. 2010;121(8). 10.1016/j.clinph.2010.02.159.

33. Verzé L, et al. Distribution of GAP-43 nerve fibers in the skin of the adult human hand. Anat Rec A Discov Mol Cell Evol Biol. 2003;272A(1):467–473.

34. Bursova S, et al. Expression of growth-associated protein 43 in the skin nerve fibers of patients with type 2 diabetes mellitus. J Neurol Sci. 2012;315(1–2). 10.1016/j.jns.2011.11.038.

35. Scheytt S, et al. Increased gene expression of growth associated protein-43 in skin of patients with early-stage peripheral neuropathies. J Neurol Sci. 2015;355(1–2). 10.1016/j.jns.2015.05.044.

36. Özdağ Acarlı AN, et al. Subepidermal Schwann cell counts correlate with skin innervation - an exploratory study. Muscle Nerve. 2022;65(4):471–479.

37. Marconi A, et al. Expression and function of neurotrophins and their receptors in human melanocytes. Int J Cosmet Sci. 2006;28(4). 10.1111/j.1467-2494.2006.00321.x.

38. Wilcox NC, et al. Interactions between skin-resident dendritic and Langerhans cells and pain-sensing neurons [preprint]. Journal of Allergy and Clinical Immunology. 2024;154(1). 10.1016/j.jaci.2024.03.006.

39. Doss ALN, Smith PG. Langerhans cells regulate cutaneous innervation density and mechanical sensitivity in mouse footpad. Neurosci Lett. 2014;578. 10.1016/j.neulet.2014.06.036.

40. Caxaria S, et al. Neutrophils infiltrate sensory ganglia and mediate chronic widespread pain in fibromyalgia. Proc Natl Acad Sci U S A. 2023;120(17):e2211631120.

41. Zhang C, Yang M, Ericsson AC. Function of Macrophages in Disease: Current Understanding on Molecular Mechanisms. Front Immunol. 2021;12. 10.3389/fimmu.2021.620510.

42. Msheik Z, et al. The macrophage: a key player in the pathophysiology of peripheral neuropathies [preprint]. J Neuroinflammation. 2022;19(1). 10.1186/s12974-022-02454-6.

43. Lund H, et al. CD163+ macrophages monitor enhanced permeability at the blood-dorsal root ganglion barrier. J Exp Med. 2024;221(2). 10.1084/jem.20230675.

44. Zaba LC, et al. Normal human dermis contains distinct populations of CD11c+BDCA-1+ dendritic cells and CD163+FXIIIA+ macrophages. J Clin Invest. 2007;117(9):2517–25.

45. Liu Z, et al. Spatiotemporal single-cell roadmap of human skin wound healing. Cell Stem Cell. 2025;32(3):479–498.e8.

46. Kosek E, Ekholm J, Hansson P. Sensory dysfunction in fibromyalgia patients with implications for pathogenic mechanisms. Pain. 1996;68(2):375–383.

47. Marshall A, et al. Small fibre pathology, small fibre symptoms and pain in fibromyalgia syndrome. Sci Rep. 2024;14(1). 10.1038/s41598-024-54365-6.

48. Sommer C, Üçeyler N. Small fiber pathology in fibromyalgia syndrome. Pain Rep. 2025;10(1):e1220.

49. Berglund B, et al. Quantitative and qualitative perceptual analysis of cold dysesthesia and hyperalgesia in fibromyalgia. Pain. 2002;96(1–2). 10.1016/S0304-3959(01)00443-2.

50. Augière T, et al. Tactile Detection in Fibromyalgia: A Systematic Review and a Meta-Analysis. *Frontiers in pain research (Lausanne*, Switzerland). 2021;2:740897.

51. Jänsch S, et al. Distinguishing fibromyalgia syndrome from small fiber neuropathy: A clinical guide. Pain Rep. 2024;9(1). 10.1097/PR9.0000000000001136.

52. Cohen SP, Vase L, Hooten WM. Chronic pain: an update on burden, best practices, and new advances [preprint]. The Lancet. 2021;397(10289). 10.1016/S0140-6736(21)00393-7.

53. Engen DJ, et al. Effects of transdermal magnesium chloride on quality of life for patients with fibromyalgia: a feasibility study. J Integr Med. 2015;13(5):306–13.

54. Lauria G, et al. EFNS guidelines on the use of skin biopsy in the diagnosis of peripheral neuropathy. Eur J Neurol. 2005;12(10):747–58.

55. Lauria G, et al. Tubule and neurofilament immunoreactivity in human hairy skin: Markers for intraepidermal nerve fibers. Muscle Nerve. 2004;30(3). 10.1002/mus.20098.

56. Castañeda-Corral G, et al. The majority of myelinated and unmyelinated sensory nerve fibers that innervate bone express the tropomyosin receptor kinase A. Neuroscience. 2011;178:196–207.

57. Evdokimov D, et al. Pain-associated Mediators and Axon Pathfinders in Fibromyalgia Skin Cells. J Rheumatol. 2020;47(1):140–148.

58. Albrecht PJ, et al. Excessive Peptidergic Sensory Innervation of Cutaneous Arteriole-Venule Shunts (AVS) in the Palmar Glabrous Skin of Fibromyalgia Patients: Implications for Widespread Deep Tissue Pain and Fatigue. Pain Medicine. 2013;14(6):895–915.

59. Al-Allaf AW, et al. Investigation of cutaneous microvascular activity and flare response in patients with fibromyalgia syndrome. Rheumatology. 2001;40(10). 10.1093/rheumatology/40.10.1097.

60. Aloisi AM, Casini I. Fibromyalgia: Chronic Pain Due to a Blood Dysfunction? Int J Mol Sci. 2025;26(9). 10.3390/ijms26094153.

61. Cowan CW, et al. Vav family GEFs link activated Ephs to endocytosis and axon guidance. Neuron. 2005;46(2):205–17.

62. Pasquale EB. Eph receptor signalling casts a wide net on cell behaviour. Nat Rev Mol Cell Biol. 2005;6(6):462–75.

63. Abdo H, et al. Specialized cutaneous Schwann cells initiate pain sensation. Science. 2019;365(6454):695–699.

64. Zhou Y, et al. Schwann Cell–Secreted S100B Promotes Wound Healing via Paracrine Modulation. J Dent Res. 2025;104(3):330–340.

65. Direder M, et al. Schwann cells contribute to keloid formation. Matrix Biology. 2022;108. 10.1016/j.matbio.2022.03.001.

66. Khudhair OS, Gorial FI, Hameed AF. Fibromyalgia Syndrome and Vitiligo: A Novel Association. Arch Rheumatol. 2018;33(2):174–180.

67. Carter MC, Metcalfe DD, Komarow HD. Mastocytosis. Immunol Allergy Clin North Am. 2014;34(1):181–96.

68. Keith YH, et al. Infiltration and local differentiation of bone marrow-derived integrinβ7-positive mast cell progenitors in atopic dermatitis-like skin. J Allergy Clin Immunol. 2023;151(1):159–171.e8.

69. Eneström S, Bengtsson A, Frödin T. Dermal IgG deposits and increase of mast cells in patients with fibromyalgia--relevant findings or epiphenomena? Scand J Rheumatol. 1997;26(4):308–13.

70. Ang DC, Hilligoss J, Stump T. Mast Cell Stabilizer (Ketotifen) in Fibromyalgia: Phase 1 Randomized Controlled Clinical Trial. Clin J Pain. 2015;31(9):836–842.

71. Coffelt SB, Wellenstein MD, de Visser KE. Neutrophils in cancer: neutral no more. Nat Rev Cancer. 2016;16(7):431–46.

72. Kaplan MJ. Role of neutrophils in systemic autoimmune diseases. Arthritis Res Ther. 2013;15(5):219.

73. Mócsai A. Diverse novel functions of neutrophils in immunity, inflammation, and beyond. J Exp Med. 2013;210(7):1283–99.

74. Villanueva E, et al. Netting neutrophils induce endothelial damage, infiltrate tissues, and expose immunostimulatory molecules in systemic lupus erythematosus. J Immunol. 2011;187(1):538–52.

75. Vicanolo T, et al. Matrix-producing neutrophils populate and shield the skin. Nature. 2025;641(8063):740–748.

76. Banfi G, et al. T Cell Subpopulations in the Physiopathology of Fibromyalgia: Evidence and Perspectives. Int J Mol Sci. 2020;21(4). 10.3390/ijms21041186.

77. Gaudet AD, Popovich PG, Ramer MS. Wallerian degeneration: gaining perspective on inflammatory events after peripheral nerve injury. J Neuroinflammation. 2011;8:110.

78. Lee KM, et al. Macrophage Function Disorders. Encyclopedia of Life Sciences. Wiley; 2013.

79. Ferreira DW, et al. CD163 overexpression using a macrophage-directed gene therapy approach improves wound healing in ex vivo and in vivo human skin models. Immunobiology. 2020;225(1). 10.1016/j.imbio.2019.10.011.

80. Kasraie S, Werfel T. Role of macrophages in the pathogenesis of atopic dermatitis. Mediators Inflamm. 2013;2013:942375.

81. Chong BF, et al. A subset of CD163+ macrophages displays mixed polarizations in discoid lupus skin. Arthritis Res Ther. 2015;17(1):324.

82. Sugaya M, et al. Association of the numbers of CD163(+) cells in lesional skin and serum levels of soluble CD163 with disease progression of cutaneous T cell lymphoma. J Dermatol Sci. 2012;68(1):45–51.

83. Wynn TA, Vannella KM. Macrophages in Tissue Repair, Regeneration, and Fibrosis. Immunity. 2016;44(3):450–462.

84. Payet CA, et al. Central Role of Macrophages and Nucleic Acid Release in Myasthenia Gravis Thymus. Ann Neurol. 2023;93(4):643–654.

85. Sim SL, et al. Macrophages in Skin Wounds: Functions and Therapeutic Potential. Biomolecules. 2022;12(11). 10.3390/biom12111659.

86. McWhorter FY, et al. Modulation of macrophage phenotype by cell shape. Proc Natl Acad Sci U S A. 2013;110(43). 10.1073/pnas.1308887110.

87. Kesapragada M, et al. A data-driven approach to establishing cell motility patterns as predictors of macrophage subtypes and their relation to cell morphology. PLoS One. 2024;19(12):e0315023.

88. Thomas SJ, et al. Abnormal Intraepidermal Nerve Fiber Density in Disease: A Scoping Review. medRxiv. [published online ahead of print: February 8, 2023]. 10.1101/2023.02.08.23285644.

89. Sierra-Silvestre E, et al. Occurrence of corneal sub-epithelial microneuromas and axonal swelling in people with diabetes with and without (painful) diabetic neuropathy. Diabetologia. 2023;66(9):1719–1734.

90. Espinoza N, Papadopoulos V. Role of Mitochondrial Dysfunction in Neuropathy. Int J Mol Sci. 2025;26(7):3195.

91. Saleh A, et al. Ciliary neurotrophic factor activates NF-κB to enhance mitochondrial bioenergetics and prevent neuropathy in sensory neurons of streptozotocin-induced diabetic rodents. Neuropharmacology. 2013;65:65–73.

92. Hakim S, et al. Macrophages protect against sensory axon loss in peripheral neuropathy. Nature. 2025;640(8057):212–220.

93. Sipka T, et al. Macrophages undergo a behavioural switch during wound healing in zebrafish. Free Radic Biol Med. 2022;192:200–212.

94. Lauria G, et al. Intraepidermal nerve fiber density at the distal leg: A worldwide normative reference study. Journal of the Peripheral Nervous System. 2010.

95. Hasegawa T, et al. Reduction in Human Epidermal Langerhans Cells with Age Is Associated with Decline in CXCL14-Mediated Recruitment of CD14+ Monocytes. Journal of Investigative Dermatology. 2020;140(7). 10.1016/j.jid.2019.11.017.

96. Ortonne JP. Pigmentary changes of the ageing skin. Br J Dermatol. 1990;122 Suppl 35:21–8.

97. Hedin P-J, et al. The Fibromyalgia Impact Questionnaire, a Swedish Translation of a New Tool for Evaluation of the Fibromyalgia Patient. Scand J Rheumatol. 1995;24(2):69–75.

98. Löfgren M, et al. Long-term, health-enhancing physical activity is associated with reduction of pain but not pain sensitivity or improved exercise-induced hypoalgesia in persons with rheumatoid arthritis. Arthritis Res Ther. 2018;20(1). 10.1186/s13075-018-1758-x.

99. Borg G, Ljunggren G, Ceci R. The increase of perceived exertion, aches and pain in the legs, heart rate and blood lactate during exercise on a bicycle ergometer. Eur J Appl Physiol Occup Physiol. 1985;54(4). 10.1007/BF02337176.

100. Pincivero DM, Coelho AJ, Erikson WH. Perceived exertion during isometric quadriceps contraction: A comparison between men and women. Journal of Sports Medicine and Physical Fitness. 2000;40(4).

101. Kosek E, Ekholm J, Nordemar R. A comparison of pressure pain thresholds in different tissues and body regions. Long-term reliability of pressure algometry in healthy volunteers. Scand J Rehabil Med. 1993;25(3):117–24.

102. Hunt MA, et al. DRGquant: A new modular AI-based pipeline for 3D analysis of the DRG. J Neurosci Methods. 2022;371:109497.

103. Ronneberger O, Fischer P, Brox T. U-Net: Convolutional Networks for Biomedical Image Segmentation. 2015.

104. Schindelin J, et al. Fiji: an open-source platform for biological-image analysis. Nat Methods. 2012;9(7):676–682.

105. Haase R, et al. CLIJ: GPU-accelerated image processing for everyone. Nat Methods. 2020;17(1):5–6.

