## Supplementary table 1. Antibodies and conditions used. for "Exploring the neuroimmune cellular landscape in the skin of subjects with fibromyalgia"

**SUPLEMENTARY MATERIAL**

Supplementary table 1. Antibodies and conditions used.

| **Target** | **Fluorophore** | **Host** | **Manufacturer** | **Catalogue #** | **Dilution** |
| --- | --- | --- | --- | --- | --- |
| **TUBB3** | **N/A** | **Mouse** | **Promega** | **G712A** | **1:500** |
| **MelanA** | **N/A** | **Sheep** | **R&D** | **AF8008** | **1:500** |
| **PGP9.5** | N/A | Mouse | Bio-Rad | 7863-1004 | 1:400 |
| **NF200** | N/A | Chicken | Neuromics | CH22104 | 1:2000 |
| **GAP43** | N/A | Rabbit | Abcam | Ab12274 | 1:200 |
| **CD31** | N/A | Sheep | R&D systems | AF806 | 1:400 |
| **CD207** | N/A | Rat | Novus Biologicals | DDX0362P | 1:1000 |
| **S100b** | N/A | Rabbit | Abcam | ab52642 | 1:500 |
| **CD68** | N/A | Mouse | DAKO | M0814 | 1:500 |
| **CD163** | N/A | Mouse | Novus Biologicals | Nb110-40686 | 1:500 |
| **CD1a** | N/A | Mouse | Bio-Rad | MCA80 | 1:1000 |
| **FcERI** | PE | Mouse | Thermofisher | 12-5899-42 | 1:200 |
| **CD117 clone 104D2** | APC | Mouse | BD | 333233 | 1:200 |
| **CD3** | N/A | Rabbit | DAKO | M7193 | 1:200 |
| **Neutrophil elastase** | N/A | Rabbit | Abcam | Ab131260 | 1:1000 |
| **Mouse IgG** | Cy2 | Donkey | Jackson Immunoresearch | 715-225-151 | 1:200 |
| **Mouse IgG** | Cy3 | Donkey | Jackson Immunoresearch | 715-165-151 | 1:600 |
| **Mouse IgG** | Cy5 | Donkey | Jackson Immunoresearch | 715-175-151 | 1:500 |
| **Rat IgG** | Cy5 | Donkey | Jackson Immunoresearch | 712-175-153 | 1:500 |
| **Chicken IgY** | Cy5 | Donkey | Jackson Immunoresearch | 703-175-155 | 1:500 |
| **Rabbit IgG** | Cy2 | Donkey | Jackson Immunoresearch | 711-225-152 | 1:200 |
| **Rabbit IgG** | Cy3 | Donkey | Jackson Immunoresearch | 711-165-152 | 1:500 |
| **Sheep IgG** | Alexa Fluor^TM^ 488 | Donkey | Invitrogen | A-11015 | 1:500 |
